# T cell activation, highly armed cytotoxic cells and a sharp shift in monocytes CD300 receptors expression is characteristic of patients with severe COVID-19

**DOI:** 10.1101/2020.12.22.423917

**Authors:** Olatz Zenarruzabeitia, Gabirel Astarloa-Pando, Iñigo Terrén, Ane Orrantia, Raquel Pérez-Garay, Iratxe Seijas-Betolaza, Javier Nieto-Arana, Natale Imaz-Ayo, Silvia Pérez-Fernández, Eunate Arana-Arri, Francisco Borrego

## Abstract

COVID-19 manifests with a wide diversity of clinical phenotypes characterized by dysfunctional and exaggerated host immune responses. Many results have been described on the status of the immune system of patients infected with SARS-CoV-2, but there are still aspects that have not been fully characterized. In this study, we have analyzed a cohort of patients with mild, moderate and severe disease. We performed flow cytometric studies and correlated the data with the clinical features and clinical laboratory values of patients. Both conventional and unsupervised data analyses concluded that patients with severe disease are characterized, among others, by a higher state of activation in all T cell subsets, higher expression of perforin and granzyme B in cytotoxic cells, expansion of adaptive NK cells and the accumulation of activated and immature dysfunctional monocytes which are identified by a low expression of HLA-DR and an intriguing abrupt change in the expression pattern of CD300 receptors. More importantly, correlation analysis showed a strong association between the alterations in the immune cells and the clinical signs of severity. These results indicate that patients with severe COVID-19 have a broad perturbation of their immune system, and they will help to understand the immunopathogenesis of severe COVID-19 as well as could be of special value for physicians to decide which specific therapeutic options are most effective for their patients.

## INTRODUCTION

Severe acute respiratory syndrome coronavirus 2 (SARS-CoV-2) can cause coronavirus disease 2019 (COVID-19) which, in the worst cases scenario, can lead to severe manifestations such as acute respiratory distress syndrome, characterized by aggressive inflammatory responses in the lower part of respiratory tract, and multiple organ failure. A relevant number of symptomatic patients require hospitalization and a portion of them are admitted to the intensive care unit (ICU), moreover death may occur in a significant number of cases (Huang et al., 2020; Zhou et al., 2020a). The thrombotic complications associated with COVID-19 represent a very important problem. Embolism and thrombosis are frequent clinical features of patients with severe COVID-19 (Klok et al., 2020; Lodigiani et al., 2020), sometimes despite anticoagulation therapy. Patients with severe disease have abnormal coagulation characteristics, including elevated D-dimer levels, and generalized thrombotic microvascular injury (Rapkiewicz et al., 2020; Tang et al., 2020; Zhou et al., 2020a).

In acute respiratory viral infections, pathology can be caused directly by the virus and/or by a damaging immune response from the host (Blanco-Melo et al., 2020; Moore and June, 2020; Vabret et al., 2020). In this sense, severe COVID-19 is due not only to the direct effects of SARS-CoV-2, but also to a misdirected host response with complex immune dysregulation (Giamarellos-Bourboulis et al., 2020; Kuri-Cervantes et al., 2020; Laing et al., 2020; Mathew et al., 2020; Su et al., 2020; Zhou et al., 2020b). Therefore, it is very important to exactly recognize and identify the immunological signatures that correlate with the severity of the disease, since this aspect undoubtedly has relevant clinical implications related to patients’ stratification and management. From the first publications, the knowledge about the dysfunctional immune response in COVID-19 is constantly evolving. Most reports on immune dysfunction in COVID-19 patients have focused on severe disease. Hence, patients with severe COVID-19 exhibit in plasma higher amounts of numerous cytokines and chemokines than less severe cases (Herold et al., 2020; Del Valle et al., 2020; Yang et al., 2020). Severe manifestations are caused, in part, by high levels of interleukin (IL)-6 and the subsequent cytokine storm together with an altered type I interferon (IFN) response with low IFN production and an altered expression of IFN-regulated genes (Hadjadj et al., 2020; Del Valle et al., 2020). The cytokine storm is characterized by systemic inflammation, hemodynamic instability, hyperferritinemia and multiple organ failure (Moore and June, 2020). Many studies of circulating immune cells by flow and mass cytometry and/or single-cell RNA sequencing have provided valuable insights into immune perturbations in COVID-19 (Carissimo et al., 2020; Giamarellos-Bourboulis et al., 2020; Kuri-Cervantes et al., 2020; Laing et al., 2020; Mathew et al., 2020; Sánchez-Cerrillo et al., 2020; Schulte-Schrepping et al., 2020; Silvin et al., 2020; Su et al., 2020; Wilk et al., 2020). Recently, a multi-omics approach study has identified a major shift between mild and moderate disease, in which increased inflammatory signaling correlates with clinical metrics of blood clotting and plasma composition changes, suggesting that moderate disease may be the most effective situation for therapeutic intervention (Su et al., 2020).

Lymphopenia, including T and NK cell lymphopenia, is a characteristic of severe COVID-19 (Giamarellos-Bourboulis et al., 2020; Zhou et al., 2020a). In addition, alterations in the T cell compartment of COVID-19 patients have been described (Kuri-Cervantes et al., 2020; Laing et al., 2020; Mathew et al., 2020; Zhou et al., 2020b). Among these, an increment in the frequency of activated and proliferating memory CD4 T cells and memory CD8 T cells in subsets of patients have been documented (Mathew et al., 2020). Also, T cell exhaustion and increased expression of inhibitory receptors on peripheral T cells have been described (Li et al., 2020; Mathew et al., 2020; Zheng et al., 2020). Nonetheless, it is important to consider that these inhibitory receptors are also increased after T cell activation (Mathew et al., 2020). Besides the evidences of T cell activation in COVID-19 patients, some studies have found decreases in polyfunctionality or cytotoxicity (Zheng et al., 2020). Other reported alterations in the T cell compartment include, for example, a decrease in γδ T cells (Laing et al., 2020). Related to B cells, it has been described an increase in the circulating plasmablasts and in proliferating B cell subsets, among others (Mathew et al., 2020). Alterations in natural killer (NK) cells during acute SARS-CoV-2 infection have also been reported. For example, reduced NK cell counts in patients with severe COVID-19 and impaired degranulating activity and IFN-gamma production in response to classical targets, such as K562 cells, have been published (Mazzoni et al., 2020; Osman et al., 2020). Other studies have shown that there is an increase in the frequency of NK cells displaying inhibitory receptors, such as NKG2A (Demaria et al., 2020; Li et al., 2020; Zheng et al., 2020). However, others have described a strong activation of both circulating and lung NK cells and an expansion of adaptive NK cells in patients with severe disease (Jiang et al., 2020; Maucourant et al., 2020).

Diverse immune mechanisms are on place to detect viral replication and protect the host. Pattern recognition receptors of the innate immune system recognize viral antigens and virus-induced damage, increasing bone marrow hematopoiesis, the release of myeloid cells including neutrophils and monocytes, and the secretion of cytokines and chemokines (Stegelmeier et al., 2019). If the inflammatory condition is not controlled, then emergency hematopoiesis may lead to bystander tissue damage that with the cytokine storm causes organ dysfunction. It is well known that the myeloid compartment is also profoundly altered during SARS-CoV-2 infection, especially in patients with severe COVID-19 (Mann et al., 2020; Schulte-Schrepping et al., 2020; Silvin et al., 2020). For example, dendritic cells (DCs) were found to be reduced in number and functionally impaired and the ratio of conventional DCs (cDCs) to plasmacytoid DCs (pDCs) was increased in patients with severe disease (Zhou et al., 2020b). Some authors have found that pDCs and CD141+ cDCs were equally diminished in patients irrespective of the disease severity, while the decrease in CD1c+ cDCs was more evident in patients with severe COVID-19, suggesting that this specific cDC subset migrates to the lungs and other locations (Sánchez-Cerrillo et al., 2020). Regarding circulating monocytes, alterations in the frequency of certain subpopulations have been described, such as transitional (CD14++CD16+) and non-classical (CD14+CD16++) monocytes (Sánchez-Cerrillo et al., 2020; Schulte-Schrepping et al., 2020; Silvin et al., 2020). Some have described that loss of non-classical monocytes could help in the identification of high risk of severe COVID-19 (Silvin et al., 2020). Patients with severe disease are characterized by an accumulation of dysfunctional activated monocytes that express low levels of HLA-DR and immature neutrophils, indicating an emergency myelopoiesis, and an accumulation of these cells in the lungs (Schulte-Schrepping et al., 2020; Silvin et al., 2020; Vitte et al., 2020). Severe COVID-19 is also characterized by a profound alteration of neutrophil subsets. Neutrophilia with immature (CD10^low^CD101-) neutrophils, indicative of emergency granulopoiesis, and dysfunctional granulocytes are characteristics of patients with severe disease (Silvin et al., 2020). A granulocytic signature has been proposed to identify SARS-CoV-2 infected from non-infected people as well as between severity stages (Vitte et al., 2020). Severity was correlated with the expression of PD-L1 in granulocytes from patients with severe COVID-19 (Schulte-Schrepping et al., 2020; Vitte et al., 2020).

Besides all the published data, there are still aspects not fully characterized in COVID-19 immunopathogenesis. Patients with severe disease exhibit a significant immune dysregulation and the nature of it is not completely understood. An in depth and complete knowledge of the dysregulated immune response is very important not only for its therapeutic implications, but also to better understand the immunopathology of the disease. Therefore, it is essential to entirely define the immune response characteristics related to disease features and determine at which stage of the disease specific therapeutic options may be most effective.

We have characterized lymphocytes (T, B and NK cells) and monocytes of patients with mild, moderate and severe disease using flow cytometry-based studies and correlated the results with clinical features and laboratory data. Comprehensive conventional and unsupervised analyses of the results showed that, in addition to others, the activation status of T lymphocytes and an increase in the cytotoxic potential of T and NK cells are correlated with the degree of the disease severity. Furthermore, we also describe an alteration in the expression of CD300 molecules in monocytes and granulocytes that, to our knowledge, was previously unrecognized. This alteration is characterized by an abrupt change in the expression of this family of receptors between patients with moderate and severe COVID-19. Altogether, our results may help physicians to make therapeutic decisions regarding the management of patients with moderate and severe COVID-19.

## RESULTS

### SARS-CoV-2 infection: Study design, clinical cohort and clinical data

Our aim was to evaluate the impact of acute SARS-CoV-2 infection in circulating leukocytes. To this end, we performed a cross-sectional study. Forty four patients with COVID-19 disease were recruited for the study. To correlate laboratory findings, including frequencies and phenotype of circulating leukocytes and the severity of the disease, we stratified our cohort of COVID-19 patients into 3 groups of those showing mild (15 patients), moderate (15 patients) and severe (14 patients) disease. The demographic, clinical characteristics and clinical laboratory values are summarized in Table S1 and Table S2. Inclusion and exclusion criteria were followed to guarantee the homogeneity of the cohort, including age, gender, severity of the disease and time from the onset of symptoms to sample collection. In addition, twelve healthy controls (HC) were included in the study.

No significant differences were found between COVID-19 patients and HC in relation to age (median ages of 64 and 59.5, respectively). There were also no significant differences between the three groups of patients (severe, moderate and mild) in relation to the number of days from the appearance of symptoms and the sample collection: median of 8 days for the mild group (range: 0 to 35), 3 days for the moderate group (range: 0 to 15) and 7 days for the severe group (range: 0 to 21). Regarding the gender of the participants, 7 (58.33%) men and 5 (41.66%) women participated in the HC group and 20 men (45.45%) and 24 (54.54%) women in the COVID-19 group (Fig. 1A). As shown in Fig. 1B, and in agreement with previous studies (Huang et al., 2020; Mann et al., 2020; Del Valle et al., 2020; Zhou et al., 2020a), we observed an increase in the levels of plasma IL-6, C-reactive protein (CRP) and ferritin in COVID-19 patients in comparison with HC (Fig. 1B). Specifically, 26% of patients exhibited IL-6 levels above the normal range (>40 pg/mL). Interestingly, all HC had IL-6 levels below the limit of detection (<3 pg/mL), while 69% of patients had >3 pg/mL of IL-6. On the other hand, 66% of patients exhibited CRP levels above the normal range (>11 mg/L) and 57% of patients exhibited ferritin levels above the normal range (>300 ng/mL) (Fig. 1B). Furthermore, although white blood cell (WBC) counts were mostly normal in mild and moderate COVID-19 patients, some moderate and severe patients exhibited high WBC counts (Fig. 1C). Also, and in accordance with the literature (Hadjadj et al., 2020; Huang et al., 2020), we observed frequencies and absolute numbers of lymphocytes below the normal values, and frequencies and absolute number of neutrophils above the normal values associated with the severity of the disease (Fig. 1C). Finally, increased levels of IL-6 (>40 pg/mL), CRP (>11 mg/L), ferritin (>300 ng/mL), fibrinogen (>400 mg/dL) and D-dimer (>500 ng/mL) and lower levels of hemoglobin (<13 g/dL) were observed mostly in moderate and severe patients (Fig. 1D).

**Fig. 1.**
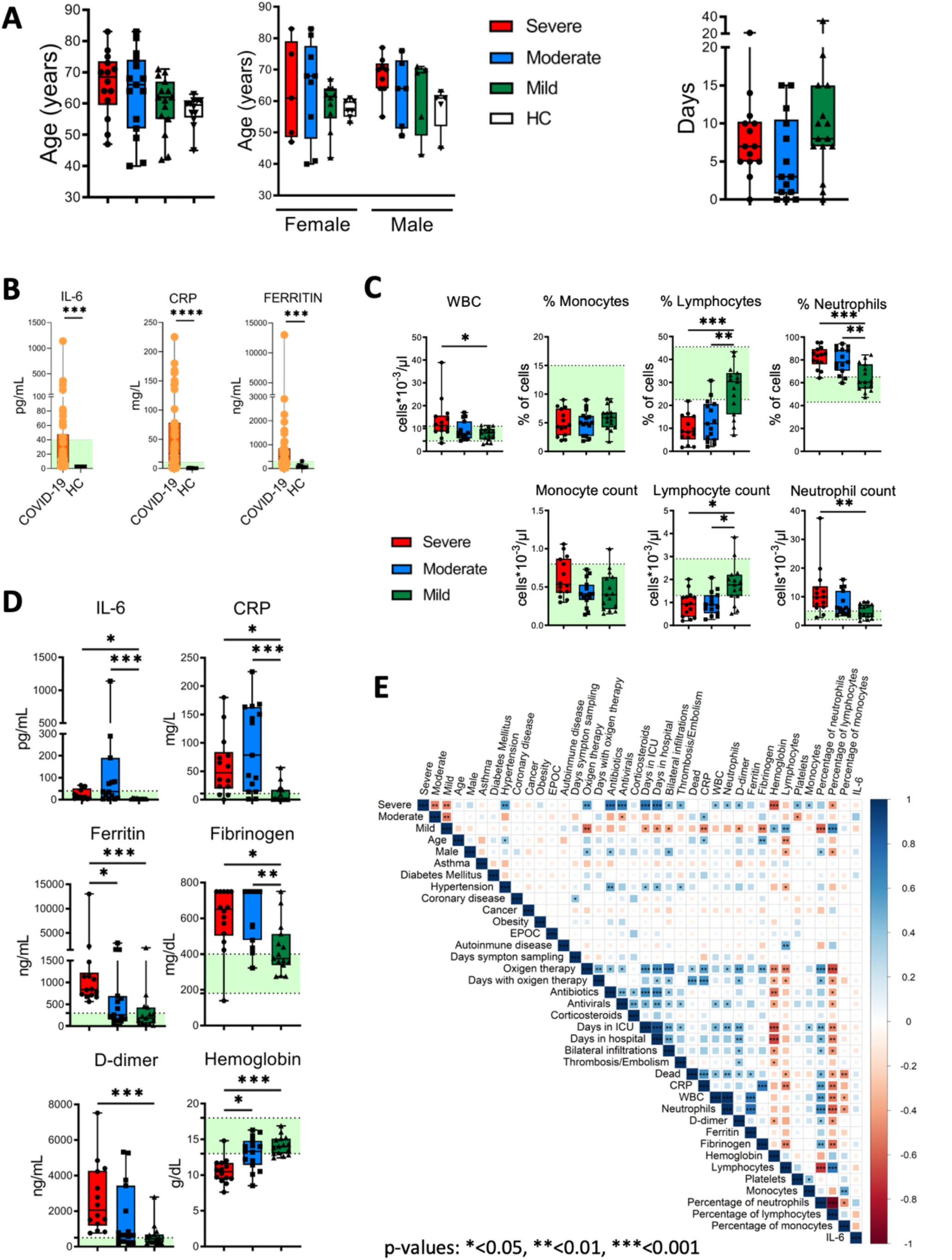
Clinical features of patients, quantification of leukocyte subsets and inflammation markers. (**A**) Left: age and gender distribution of patient cohorts in this study, including healthy controls (HC) and patients with mild (green), moderate (blue) and severe (red) COVID-19. Right: days from symptom onset to sample collection. (**B**) Plasma levels of IL-6, C reactive protein (CRP) and ferritin in HC and COVID-19 patients. The ranges of normal clinical laboratory values are represented in light green. (**C**) White blood cells (WBC) counts, leukocyte subsets frequencies and counts in patients with mild, moderate and severe COVID-19. The light green region represents the normal range for healthy people in the clinical laboratory. (**D**) Plasma levels of IL-6, CRP, ferritin, fibrinogen, D-dimer and hemoglobin in COVID-19 patients. Normal clinical laboratory values are represented in light green. (**E**) Correlogram showing Spearman correlation of the indicated clinical features for COVID-19 patients. Data in Fig. 1A, 1C and 1D are represented as boxplot graphs with the median and 25^th^ to 75^th^ percentiles, and the whiskers denote lowest and highest values. Each dot represents a donor. Significance was determined by the Kruskal-Wallis test followed by Dunn’s multiple comparison test. *p <0.05, **p <0.01, and ***p <0.001.

To examine potential associations between these general laboratory values and other clinical features, we performed correlation analysis (Fig. 1E). The analysis revealed associations between different degrees of severity with clinical features (oxygen therapy, bilateral infiltrations), comorbidities (hypertension), laboratory values (CRP, D-dimer, fibrinogen, hemoglobin), etc. Frequencies and absolute values of different subsets of WBCs were also correlated with severity degrees. Interestingly, the analysis did not reveal a correlation between IL-6 levels with other parameters. Thus, COVID-19 patients presented varied and complex clinical phenotypes and laboratory values, including evidences of inflammation and altered leukocyte counts in many patients.

### SARS-CoV-2 infection is associated with activated CD4 T cells subsets expressing higher levels of PD-1 and perforin

We next performed a detailed multiparametric flow cytometry analysis to further investigate circulating leukocytes status in COVID-19 patients (see gating strategy for each cell population in Fig. S1). Given the important role of T cells in the defense against viral infections and in the establishment of an immunological memory, as well as in the immunopathology and damage that may occur, we studied T cell subpopulations. We did not observe significant differences in the frequency of the major T cells subsets, i.e. CD4, CD8 and double negative (DN) neither in the CD4/CD8 ratio between the patients and compared with the HC (Fig. S2). Four major CD4 T cell subpopulations were examined by using the combination of CD45RA and CD27 to define naïve (CD27+CD45RA+), memory (CD27+CD45RA-), effector-memory (CD27-CD45RA-), and terminal differentiated effector-memory (TEMRA) (CD27-CD45RA+) cells (Fig. 2A). There were no significant differences in the frequencies of the four subsets between HC and COVID-19 patients. Nevertheless, the frequency of CD4 TEMRA cells was highly variable in COVID-19 patients, in which a subset of them was characterized by a relatively high number of this cell type (Fig. 2A). Most viral infections induce proliferation and activation of T cells. The latter is detected by the coexpression of CD38 and HLA-DR (Mathew et al., 2020). We found an expansion in the CD38+HLA-DR+ subset in all the non-naïve CD4 T cell subsets from COVID-19 patients, more significantly in the severe group (Fig. 2B and Fig. S3A). This expansion could be antigen-driven activation, as well as bystander activation and homeostatic proliferation. Nevertheless, the magnitude of CD38+HLA-DR+ cells expansion varied widely in our cohort and is significantly lower than the one observed in CD8 T cells (see below). After antigen recognition and activation, T cells up-regulate the expression of inhibitory receptors, such as programed cell death-1 (PD-1), with the aim of preventing an excessive response that, if not properly regulated, could be harmful to the host (Schönrich and Raftery, 2019). Therefore, in the context of an acute infection, PD-1 could also be considered an activation marker, while during chronic stimulation, T cells became progressively dysfunctional and exhausted, and the expression of PD-1 persists (Schönrich and Raftery, 2019). We studied the expression of PD-1 on CD4 T cells and found that there was an increase in all non-naïve CD4 T cells from patients, which was statistically significant in the effector-memory subset from moderate and severe COVID-19 patients (Fig. 2C and Fig. S3B). Similar to the CD38+HLA-DR+ cells, the expansion of PD-1+ cells varied widely, but importantly a strong positive correlation between CD38+HLA-DR+ cells and PD-1+ cells was observed in COVID-19 patients (Fig. 2D), suggesting the possibility that PD-1 expression on CD4 T cells during acute SARS-CoV-2 infection is more an activation marker than an exhaustion marker.

**Fig. 2.**
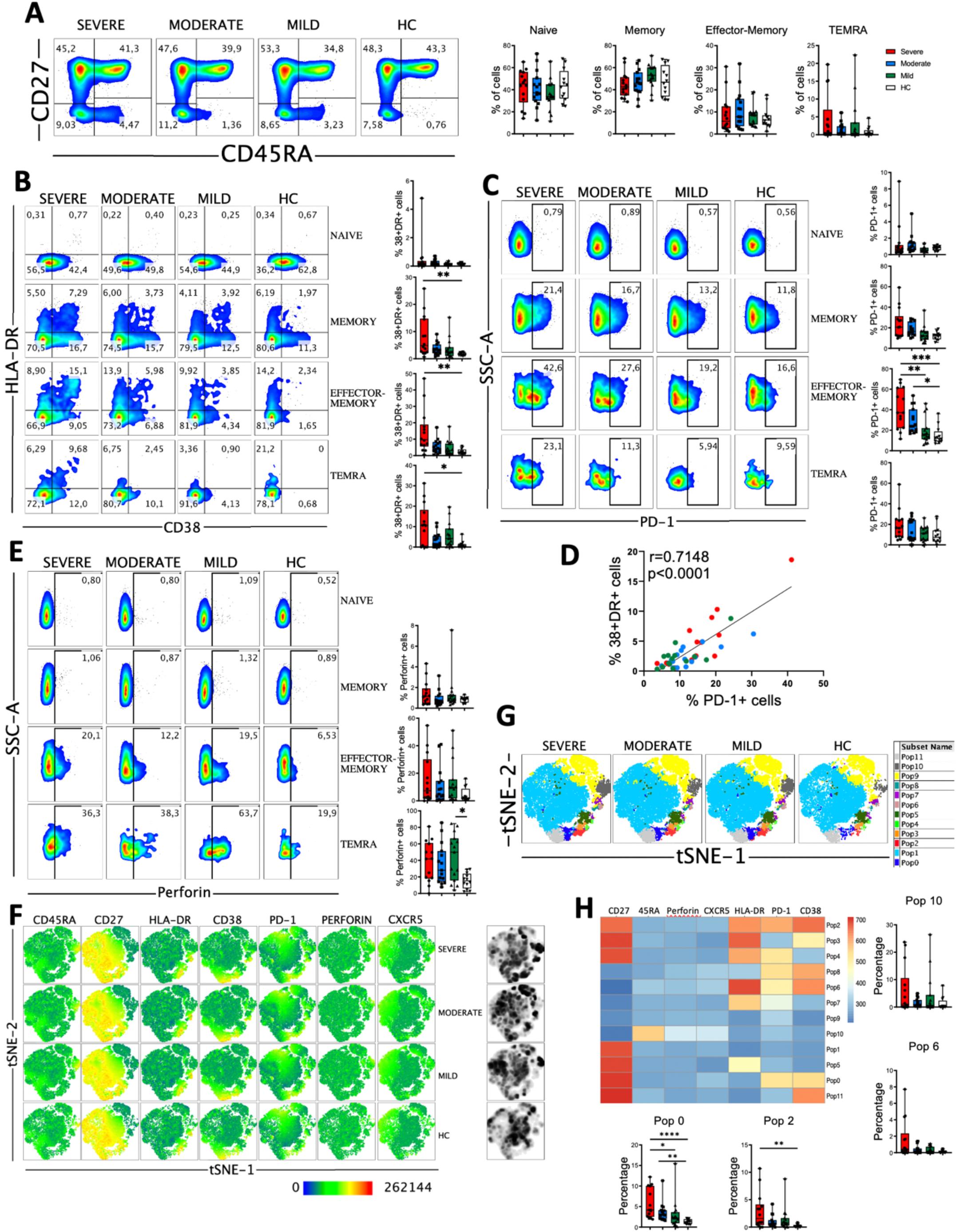
CD4 T cell subsets, activation status and perforin expression in COVID-19 patients. (**A**) Left: pseudocolor plots of concatenated peripheral CD4 T cells from healthy controls (HC) and patients with mild, moderate and severe disease. Four cell subsets were identified: naïve (CD27+CD45RA+), memory (CD27+CD45RA-), effector-memory (CD27-CD45RA-), and terminal differentiated effector-memory (TEMRA) (CD27-CD45RA+). Numbers in the quadrants are the average of each subset. Right: boxplot graphs representation of the data. (**B**). Pseudocolor plots of concatenated peripheral CD4 T cells from HC and COVID-19 patients and boxplot graphs of the frequencies of activated naïve, memory, effector-memory and TEMRA cells. Numbers in the quadrants are the average of each subset. Activated T cells are identified by the coexpression of CD38 and HLA-DR. (**C**) Pseudocolor plots of concatenated peripheral CD4 T cells and boxplot graphs showing the frequencies of PD-1+ naïve, memory, effector-memory and TEMRA cells. Numbers in the gates are the average of PD-1+ cells in each subset. (**D**) Spearman correlation of activated (CD38+HLA-DR+) with PD-1+ CD4 T cells from patients with mild, moderate and severe COVID-19. (**E**) Pseudocolor plots of concatenated peripheral CD4 T cells and boxplot graphs of the frequencies of perforin positive naïve, memory, effector-memory and TEMRA cells. Numbers in the gates are the average of perforin positive cells in each subset. (**F**) tSNE projection of the indicated markers and density plots in non-naïve CD4 T cells for all HC and COVID-19 patients. (**G**) tSNE projection of non-naïve CD4 T cell populations (Pop) identified by FlowSOM clustering tool. (**H**) Fluorescence intensity of each Pop as indicated in the column-scaled z-score and boxplot graphs showing the frequencies of Pop0, Pop2, Pop6 and Pop10 in HC and COVID-19 patients. Boxplots show the median and 25^th^ to 75^th^ percentiles, and the whiskers denote lowest and highest values. Each dot represents a donor. Significance of data in Fig. 2A, 2B, 2C, 2E and 2H was determined by the Kruskal-Wallis test followed by Dunn’s multiple comparison test. *p <0.05, **p <0.01, and ***p <0.001.

Given the relevant role of antibodies in the response to SARS-CoV-2, we also analyzed the circulating T follicular helper (TFH) cells (PD-1+CXCR5+) (Crotty, 2019). We did not observe a significant increase, except for the moderate group of patients, in the frequency of TFH in COVID-19 patients (Fig. S3C). Nevertheless, the frequency of CD38+HLA-DR+ TFH cells was expanded in patients, suggesting that they had a recent antigen encounter and have emigrated from the germinal center (Crotty, 2019) (Fig. S3C). In addition, we performed an analysis of B cells and, while the frequencies of CD27-B cells, which include mostly the naïve subset, tended to increase in COVID-19 patients, the frequencies of CD27+ memory B cells tended to decrease with the disease severity, although not significantly (Fig. S4A). Contrarily, the frequency of plasmablasts (CD27+CD38+) increased, except in patients with severe disease (Fig. S4A). We also observed a significant decrease in CXCR5 (Fig. S4B) and HLA-DR (Fig. S4C) expression levels in B cells from COVID-19 patients.

Cytotoxic CD4 T cells represent an additional mechanism by which CD4 T cells contribute to immunity. In viral infections, these perforin expressing CD4 T cells have been shown to play a protective and/or pathologic role (Broadley et al., 2017; Sanchez-Martinez et al., 2019). Therefore, we measured the expression of perforin in CD4 T cells from COVID-19 patients (Fig. 2E and Fig. S3D). Results showed that the frequency of effector-memory and TEMRA CD4 T cells expressing perforin from a subset of COVID-19 patients was higher than in HC (Fig. 2E). Although the increase in the frequency of the memory cells that express perforin was not statistically significant between COVID-19 patients and HC (Fig. 2E), we observed an enhanced perforin expression per cell basis as shown by an increase in the median fluorescence intensity (MFI) of perforin+ cells (Fig. S3D). Altogether, these results suggest that cytotoxic CD4 T cells may contribute to the clinical course of those patients.

To gain more insight, we performed high-dimensional mapping of seven parameters flow cytometry data in non-naïve CD4 T cells. For that, a t-distributed stochastic neighbor embedding (tSNE) representation (heatmap and density plot) of the data highlighted some regions of non-naïve CD4 T cells that were preferentially found in COVID-19 patients (Fig. 2F). Among these, cells expressing CD45RA and perforin were expanded in COVID-19 patients. To further define and also quantify these differences, we performed FlowSOM clustering and compared the expression of the seven markers to define 12 clusters or populations (or Pop) (Fig. 2G). Using this approach we identified several populations differentially expressed between HC and COVID-19 patients (Fig. 2H and Fig. 3SE). For example, Pop10, which identified TEMRA cells (CD27-CD45RA+) expressing perforin, was expanded in patients. Memory cells (CD27+CD45RA-) that express PD-1 and CD38 (Pop0) and PD-1, CD38 and HLA-DR (Pop2), as well as CD27-CD45RA^low^ expressing PD-1, CD38 and HLA-DR (Pop6) were also expanded in COVID-19 patients (Fig. 2H). Thus, COVID-19 patients were characterized by expanded populations of activated, PD-1 and perforin expressing CD4 T cells in a subgroup of patients.

### SARS-CoV-2 acute infection is associated with CD8 T cell activation in severe patients

CD8 T cells have a very relevant role in viral infections through their ability to recognize and kill virus infected cells and in the formation of the immunological memory. But also, highly differentiated CD8 T cells have been suggested to induce damage in SARS-CoV-2 infected lungs in an antigen-independent manner (AN and DW, 2020). Therefore, we next examined the four major subpopulations (naïve, memory, effector-memory and TEMRA). We observed no significant differences in the frequencies of naïve, memory and TEMRA subsets between HC and COVID-19 patients. Nevertheless, the frequency of CD8 effector-memory cells was significantly higher in patients with severe disease (Fig. 3A).

**Fig. 3.**
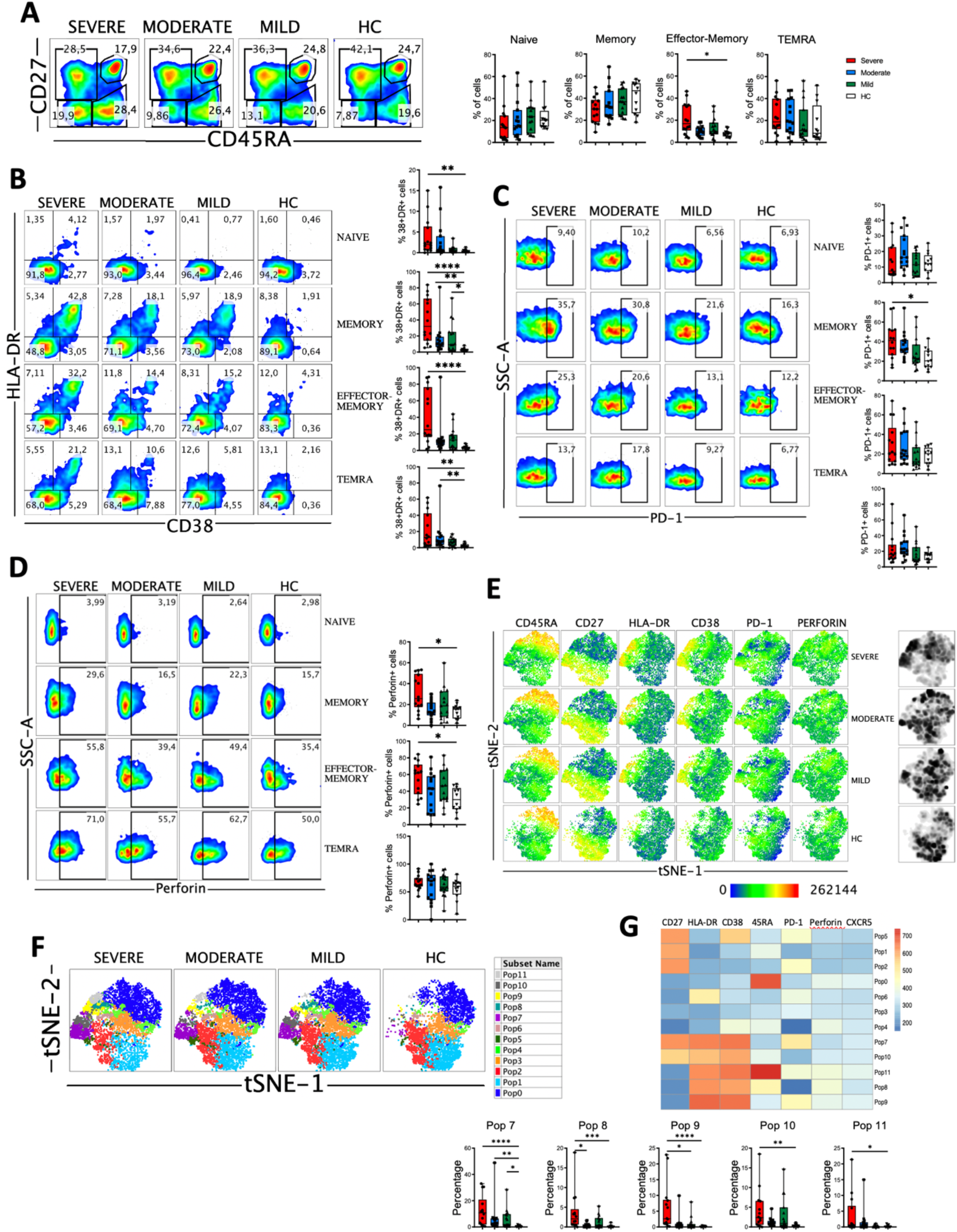
CD8 T cell subsets, perforin expression and activated cells in COVID-19 patients. (**A**) Left: pseudocolor plots of concatenated peripheral CD8 T cells from healthy controls (HC) and patients with mild, moderate and severe COVID-19. Four cell subsets were identified: naïve (CD27+CD45RA+), memory (CD27+CD45RA-), effector-memory (CD27-CD45RA-), and terminal differentiated effector-memory (TEMRA) (CD27-CD45RA+). Numbers in the gates are the average of each subset. Right: boxplot graphs representation of the data. (**B**). Pseudocolor plots of concatenated peripheral CD8 T cells from HC and COVID-19 patients and boxplot graphs of the frequencies of activated naïve, memory, effector-memory and TEMRA cells. Numbers in the quadrants are the average of each subset. Activated T cells are identified by the coexpression of CD38 and HLA-DR. (**C**) Pseudocolor plots of concatenated peripheral CD8 T cells and boxplot graphs showing the frequencies of PD-1+ naïve, memory, effector-memory and TEMRA cells. Numbers in the gates are the average of PD-1+ cells in each subset. (**D**) Pseudocolor plots of concatenated peripheral CD8 T cells and boxplot graphs of the frequencies of perforin positive naïve, memory, effector-memory and TEMRA cells. Numbers in the gates are the average of perforin positive cells in each subset. (**E**) tSNE projection of the indicated markers and density plots in non-naïve CD8 T cells for all HC and COVID-19 patients. (**F**) tSNE projection of non-naïve CD8 T cell populations (Pop) identified by FlowSOM clustering tool. (**G**) Fluorescence intensity of each Pop as indicated in the column-scaled z-score and boxplot graphs showing the frequencies of Pop7, Pop8, Pop9, Pop10 and Pop11 in HC and COVID-19 patients. Boxplots show the median and 25^th^ to 75^th^ percentiles, and the whiskers denote lowest and highest values. Each dot represents a donor. Significance of data in Fig. 3A, 3B, 3C, 3D and 3G was determined by the Kruskal-Wallis test followed by Dunn’s multiple comparison test. *p <0.05, **p <0.01, ***p <0.001, and ****p <0.0001.

Then, we determined the activation status of CD8 T cells. We observed that COVID-19 patients exhibited a significant expansion of activated (CD38+HLA-DR+) cells, especially in the patients with severe disease (Fig. S5A). As in CD4 T cells, this expansion could be not only antigen-driven activation, but also homeostatic proliferation and bystander activation. When we looked at the CD8 T cells subsets, increased frequencies of activated CD8 T cells were also observed in patients, most significantly in those with severe disease (Fig. 3B). The magnitude of the activated cells expansion varied widely, although in a significant subset of patients with severe COVID-19 more than 50% of their memory, effector-memory and TEMRA CD8 T cells were activated compared to less than 10% in all HC (Fig. 3B). We also studied the expression of PD-1 on CD8 T cells (Fig. 3C and Fig. S5B) and although there was a tendency, no significant differences were observed between HC and patients, with the exception of an expansion of PD-1+ CD8 memory cells in patients with severe disease. In contrast to the CD4 T cells, we did not observe a correlation between CD38+HLA-DR+ cells and PD-1+ cells in COVID-19 patients (Fig. S5C), probably suggesting that PD-1 expression on CD8 T cells is more a marker of exhaustion than of activation. Nevertheless, more studies are required to confirm this statement.

CD8 T cells exert their cytotoxic activity after encountering virus-infected cells by releasing perforin and granzymes that are contained in their lytic granules (Halle et al., 2017). We next determine perforin expression in our cohort of COVID-19 patients. When we looked at the total CD8 T cell population we observed a significant increase in the frequency of cells containing perforin in patients with severe disease (Fig. S5D). This could be explained by an increased in the frequency of memory and effectormemory cells expressing perforin in severe patients (Fig. 3D). Altogether, these results suggest that an expansion of activated and perforin containing non-naive CD8 T cells may contribute to the severity of the COVID-19 disease.

Projecting the non-naïve CD8 T cell subsets into the high-dimensional tSNE space (heatmap and density plot) also identified alterations in the response of these cells during SARS-CoV-2 infection compared with HC (Fig. 3E). Among others, a relevant expansion of activated non-naïve CD8 T cells was observed, more significantly in patients with severe disease. To gain more insight into the CD8 T cell alterations, we again used the FlowSOM clustering tool and compared the expression of several markers to define 12 populations (or Pop) (Fig. 3F). We were able to identify some populations that were differentially expressed between COVID-19 patients and HC (Fig. 3G and S5E). The populations containing the activated cells (Pops 7, 8, 9, 10 and 11) were significantly expanded in patients, especially in those with a severe disease. Pop7 identified activated memory CD8 T cells that are PD-1+, while Pop10 represented activated memory CD8 T cells that are PD-1-. Pop8 and Pop9 identified activated effector-memory CD8 T cells that are PD-1- and PD-1+, respectively (Fig. 3G). Pop11 identified the activated TEMRA cell subset, that was significantly increased in severe patients, while the frequency of Pop0, which identified the TEMRA non-activated cells, was the same in HC and COVID-19 patients (Fig. 3G and S5E). Pop1 identified the non-activated (CD38-HLA-DR-) and PD-1-memory subset that decreased with the severity of the disease (Fig. S5E). Hence, COVID-19 patients were characterized by an expansion of non-naïve activated CD8 T cells, including both PD-1+ and PD-1-cells.

We have also analyzed the DN T cells, and in a similar way to what happens with CD4 and CD8 T cells, the activated (HLA-DR+CD38+) and perforin expressing DN T cells are significantly expanded in COVID-19 patients (Fig. S6).

### Activated monocytes, decreased levels of HLA-DR and a shift in CD300 receptors expression pattern correlate with severe COVID-19

The myeloid cell compartment is profoundly dysregulated in patients with severe COVID-19 (Mann et al., 2020; Sánchez-Cerrillo et al., 2020; Schulte-Schrepping et al., 2020; Silvin et al., 2020; Vitte et al., 2020). Among other findings, it has been described an emergency myelopoiesis with accumulation of dysfunctional and immature monocytes that are characterized by low expression of HLA-DR and CD163 (Schulte-Schrepping et al., 2020; Silvin et al., 2020). In relation to the three main monocyte subpopulations (classical, transitional and non-classical), several authors have reported that the frequency of the CD14^low^CD16^hlgh^ (non-classical) monocyte subpopulation is decreased in patients with severe disease (Sánchez-Cerrillo et al., 2020; Silvin et al., 2020). In our cohort, we did not observe significant differences in the three main monocyte subsets, with the exception of transitional monocytes in patients with moderate disease (Fig. 4A). CD163 is a receptor expressed on monocytes that has been investigated as a potential inflammation marker in different infectious diseases (Tippett et al., 2011). We found a significant increase in the percentage of CD163+ monocytes in patients with moderate and severe disease (Fig. 4B). This increased frequency was observed in all the monocyte subsets, with a significant number of patients with moderate and severe disease exhibiting more than 40% of CD163+ transitional monocytes (Fig. 4B). Also, as previously published by others (Giamarellos-Bourboulis et al., 2020; Schulte-Schrepping et al., 2020; Silvin et al., 2020), we found a gradual decrease in HLA-DR expression levels in all monocyte subsets that correlated with the severity of the disease (Fig. 4C).

**Fig. 4.**
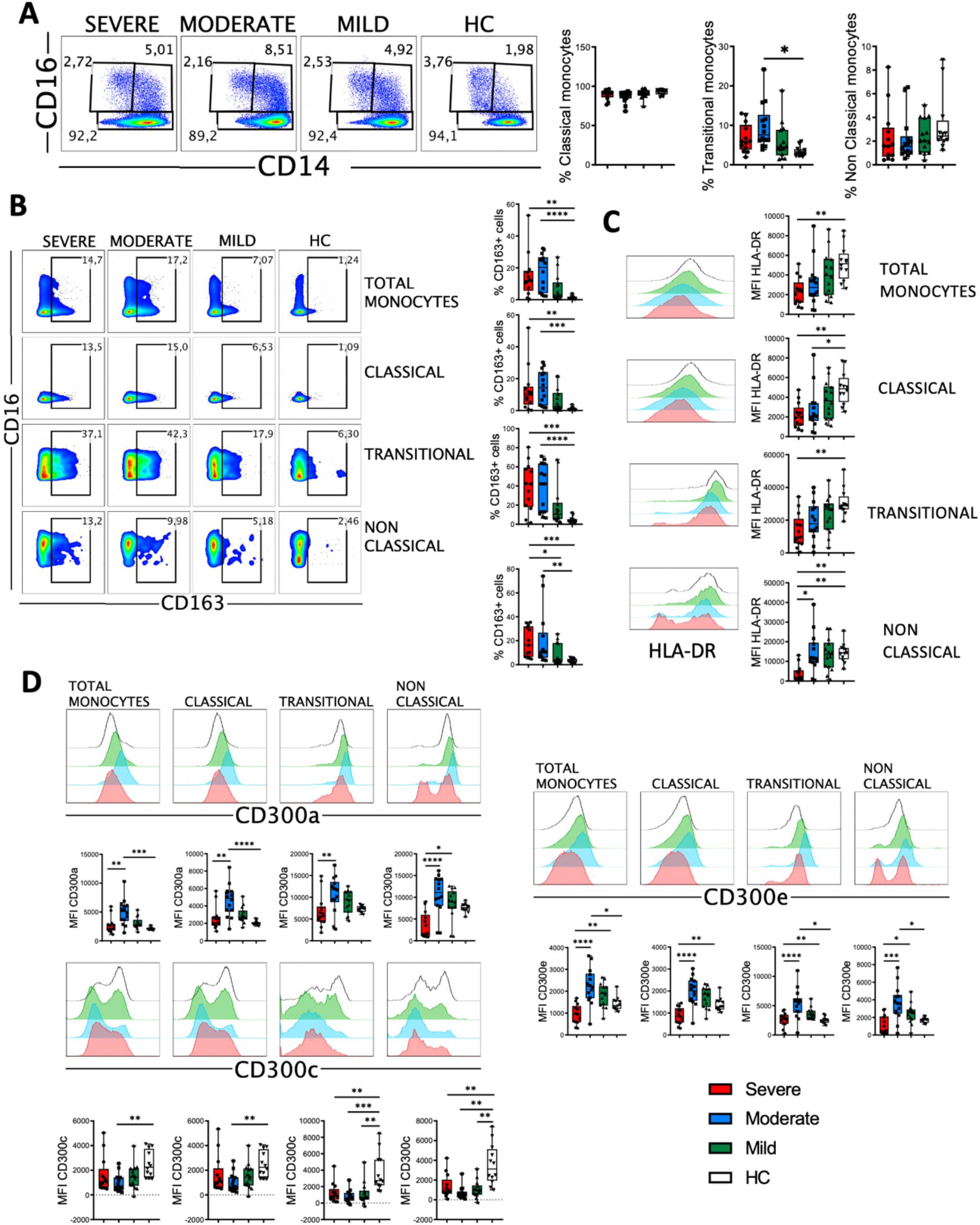
CD163, HLA-DR and CD300 receptors expression in monocytes from COVID-19 patients. (**A**) Pseudocolor plots of concatenated monocytes cells from healthy controls (HC) and patients and boxplot graphs of the frequencies of classical (CD14++CD16-), transitional (CD14++CD16+) and non-classical (CD14+CD16++) monocyte subsets. Numbers in the gates are the average of each subset. (**B**) Pseudocolor plots of all concatenated monocytes and subsets and boxplot graphs showing the frequencies of CD163+ cells. Numbers in the gates are the average of CD 163+ cells in each subset. (**C**) Histograms of concatenated monocytes and boxplot graphs showing the median fluorescence intensity (MFI) of HLA-DR in all and each monocyte subset. (**D**) Histograms of concatenated monocytes and boxplot graphs of the MFI of CD300a, CD300c and CD300e in all and each monocyte subset. Boxplots show the median and 25^th^ to 75^th^ percentiles, and the whiskers denote lowest and highest values. Each dot represents a donor. Significance of data was determined by the Kruskal-Wallis test followed by Dunn’s multiple comparison test. *p <0.05, **p <0.01, ***p <0.001, and ****p <0.0001.

The CD300 molecules are type I transmembrane proteins expressed on the surface of immune cells that modulate a multitude of signaling pathways and have been found to be involved in several diseases, including viral infections and sepsis (Borrego, 2013; Vitallé et al., 2019; Zenarruzabeitia et al., 2015, 2016). Therefore we decided to determine the expression of the inhibitory receptor CD300a and activating receptors CD300c and CD300e on monocytes from patients with COVID-19 (Fig. 4D). Results showed a differential expression of these receptors between HC and patients. Very interestingly, we observed a sharp shift in the expression of this family of receptors between moderate and severe disease. Specifically, while the expression levels of CD300a and CD300e increased in monocytes from patients with mild and moderate disease, a drastic decrease was observed in patients with a severe form of COVID-19. This decrease in the expression levels reached similar or lower levels than those expressed by patients with mild disease and HC (Fig. 4D). The expression of CD300a in granulocytes (CD66b+ cells) was similar to monocytes, with an increase in patients with mild and moderate disease and then a sharp decline in patients with severe COVID-19 (Fig. S7). The expression of CD300c on monocytes exhibited a somewhat opposite pattern to that of CD300a and CD300e (Fig. 4D). In fact, the expression of this marker decreased in all patients, and significantly in those with a moderate disease. Although the mechanism responsible for the altered expression levels of CD300 molecules in patients with COVID-19 it is not known, these results, along with other observations, may be of great importance for distinguishing those patients who have a severe disease from others who have a moderate or mild illness.

Next, we performed high-dimensional mapping of the seven parameter flow cytometry data using tSNE representation (heatmap and density plot) and we observed that some regions were preferentially found in patients when compared with HC (Fig. 5A). Then, clustering was performed using FlowSOM and 10 clusters or populations (Pop) were identified and quantified (Fig. 5B). Several populations were differentially expressed between HC and COVID-19 patients (Fig. 5C and Fig. S8). For example, the classical monocytes Pop0 and Pop4, with low expression of HLA-DR, CD300a and CD300e, were significantly expanded in patients with severe disease. Interestingly, Pop0 expressed high levels of the CD300c and Pop4 exhibited low levels of this receptor. Pop5 also belongs to the classical monocyte subset and is characterized by higher expression of HLA-DR, low CD300c and medium levels of CD300a and CD300e. In accordance with the results from conventional analysis, Pop5 was most expanded in patients with mild and moderate disease, while the frequency of this population did not change in severe COVID-19 patients when compared with HC. Pop1, which is the predominant population in HC (approximately 50% of monocytes) and is significantly reduced in patients, belongs to the classical monocyte subset and is characterized by high HLA-DR and CD300c expression, intermediate levels of CD300e and high levels of CD300a. Altogether, monocytes from COVID-19 patients have an activated phenotype, a gradual loss of HLA-DR expression that correlates with the severity of the disease and, very interestingly, a shift in the expression of CD300 molecules between moderate and severe disease.

**Fig. 5.**
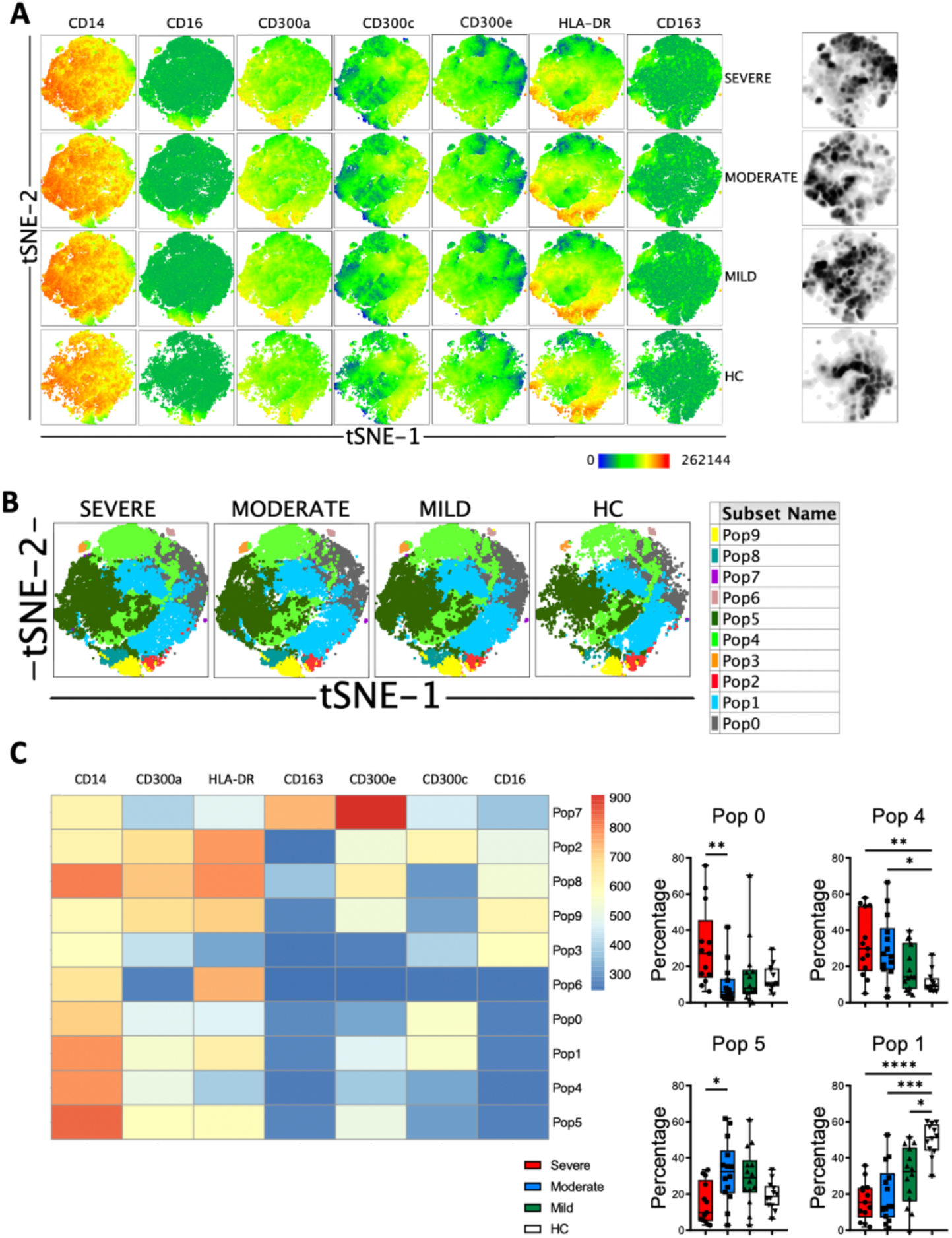
Unsupervised analysis of monocytes in COVID-19 patients. (**A**) tSNE projection of the indicated markers and density plots in monocytes for all HC and COVID-19 patients. (**B**) tSNE projection of monocyte populations (Pop) identified by FlowSOM clustering tool. (**C**) Fluorescence intensity of each Pop as indicated in the column-scaled z-score and boxplot graphs showing the frequencies of Pop0, Pop1, Pop4 and Pop5 in HC and COVID-19 patients. Boxplots show the median and 25th to 75th percentiles, and the whiskers denote lowest and highest values. Each dot represents a donor. Significance of data was determined by the Kruskal-Wallis test followed by Dunn’s multiple comparison test. *p <0.05, **p <0.01, ***p <0.001, and ****p <0.0001.

### Perforin and granzyme B armed NK cell subsets are expanded in patients with severe disease

The role of NK cells in the recognition and elimination of virus-infected cells is well documented, as well in modulating the adaptive immune response (Lam and Lanier, 2017). But also, uncontrolled NK cell activation may contribute to hyper-inflammation and tissue injury (Li et al., 2012). The exact status of NK cells in SARS-CoV-2 infection is not well elucidated and conflicting results related to their phenotype and functionality have been published. Several studies in COVID-19 patients have revealed that circulating NK cells are lower in numbers (Giamarellos-Bourboulis et al., 2020; Maucourant et al., 2020), displayed an inhibitory-related phenotype and have altered cytotoxic and immunomodulatory functions (Demaria et al., 2020; Li et al., 2020; Mazzoni et al., 2020; Osman et al., 2020; Zheng et al., 2020). Single-cell RNA sequencing studies have corroborated this trend (Wilk et al., 2020). However, other articles have described a strong activation of both, circulating and lung NK cells measured at protein and RNA levels, as well as an expansion of adaptive NK cells in patients with severe disease (Jiang et al., 2020; Maucourant et al., 2020). Therefore, we decided to perform a phenotypical analysis of NK cells from COVID-19 patients. First, we determined the frequency of the three circulating NK cell subsets, i.e. CD56^bright^, CD56^dim^ and CD56^neg^ NK cells. CD56^bright^ cells are considered a less mature subset than CD56^dim^ NK cells (Di Vito et al., 2019). Our results showed that there were no significant differences in the frequencies of the three subsets when we compared HC with COVID-19 patients (Fig. 6A).

**Fig. 6.**
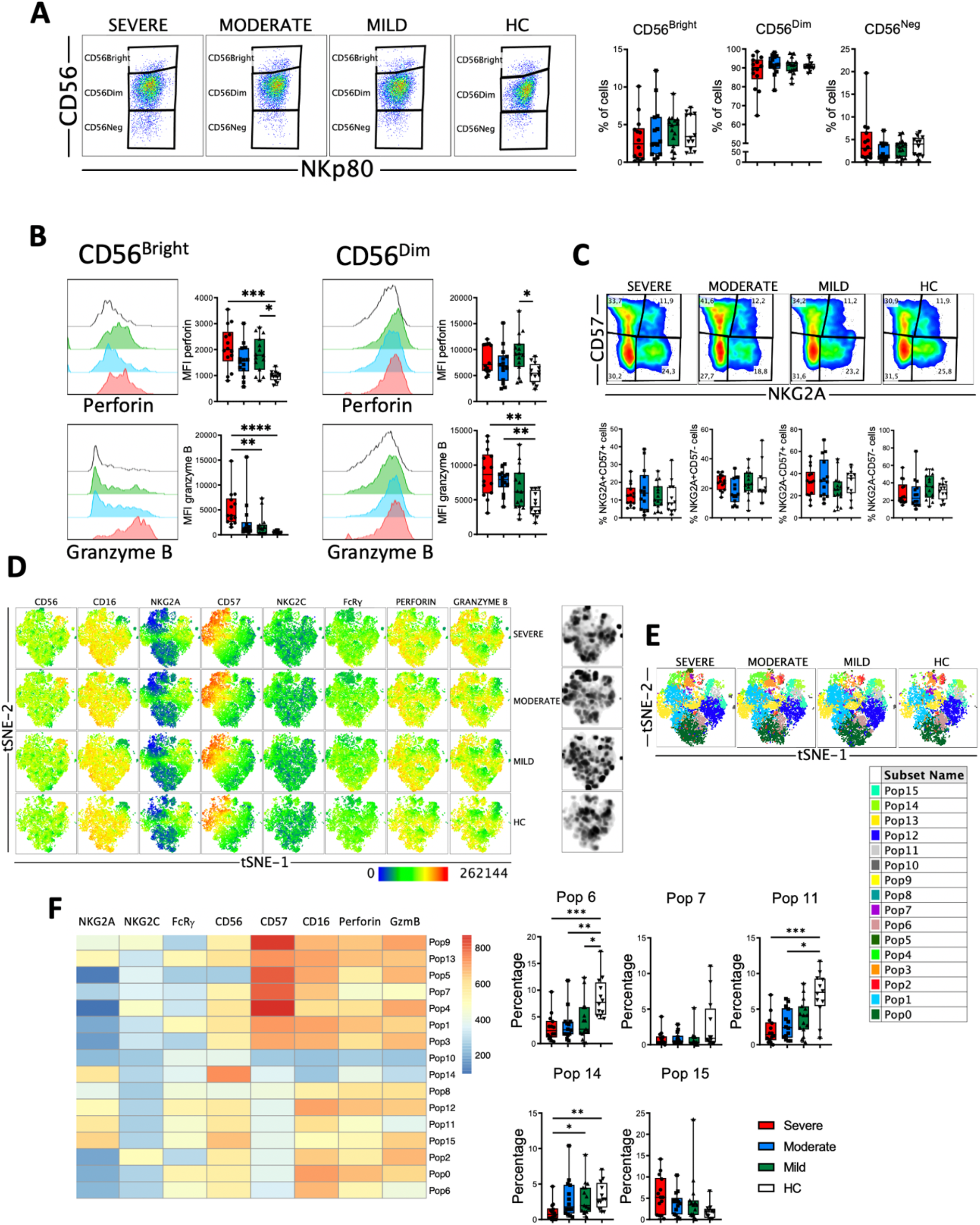
Perforin and granzyme B expression in NK cell subsets from COVID-19 patients. (**A**) Pseudocolor plots of concatenated NK cells from healthy controls (HC) and patients and boxplot graphs of the frequencies of CD56^bright^ (CD56++NKp80+), CD56^dim^ (CD56+NKp80+) and CD56^neg^ (CD56-NKp80+) NK cell subsets. (**B**) Histograms of concatenated CD56^bright^ (left) and CD56^dim^ (right) NK cells and boxplot graphs of the median fluorescence intensity (MFI) of perforin (upper) and granzyme B (lower). (**C**) Pseudocolor plots of concatenated CD56^dim^ NK cells from HC and patients and boxplot graphs of the frequencies of the four subsets based in the expression of the CD57 and NKG2A differentiation markers. Numbers in the gates are the average of each subset. (**D**) tSNE projection of the indicated markers and density plots in NK cells for all HC and COVID-19 patients. (**E**) tSNE projection of NK cells populations (Pop) identified by FlowSOM clustering. (**F**) Fluorescence intensity of each Pop as indicated in the column-scaled z-score and boxplot graphs showing the frequencies of Pop6, Pop7, Pop11, Pop14 and Pop15 in HC and COVID-19 patients. Boxplots show the median and 25^th^ to 75^th^ percentiles, and the whiskers denote lowest and highest values. Each dot represents a donor. Significance of data in Fig. 6A, 6B and 6C was determined by the Kruskal-Wallis test followed by Dunn’s multiple comparison test. *p <0.05, **p <0.01, ***p <0.001, and ****p <0.0001.

Next, we analyzed the expression of perforin and granzyme B in NK cells (Fig. 6B). Results showed that both CD56^bright^ and CD56^dim^ NK cells from COVID-19 patients exhibited higher levels of perforin and granzyme B as determined by the MFI of these markers. This increase in perforin and granzyme B levels were associated with the severity of the disease (Fig. 6B), suggesting that NK cells from patients with moderate and, even more with severe disease, have the potential to eliminate more efficiently target cells. As a subset, CD56^dim^ NK cells continue to differentiate (Di Vito et al., 2019). During this process, among other phenotypical features, they lose the expression of NKG2A and sequentially acquire CD57 (Di Vito et al., 2019). Therefore, we determined the expression of NKG2A and CD57 on CD56^dlm^ NK cells from HC and COVID-19 patients to evaluate their maturation status. Results showed that there were no significant differences in the frequencies of the four subsets between HC and patients (Fig. 6C). Nevertheless, the increased expression of perforin and granzyme B that associated with the severity of the disease was also evident in each of the four subsets (Fig. S9A).

We also performed high-dimensional mapping of the eight parameter flow cytometry data using tSNE representation and it was evident that some regions were preferentially found in COVID-19 patients when compared with HC (Fig. 6D). To gain more insight into the NK cell alterations observed in COVID-19, we used the FlowSOM clustering tool and compared the expression of the eight markers to define 16 populations (Fig. 6E). Using this approach, we were able to identify some populations that were differentially expressed between COVID-19 patients and HC (Fig. 6F and S9B). Pop6, Pop7 and Pop11 represented CD56^dim^ NK cells with relatively low amount of perforin and granzyme B expression. The frequency of these three populations was significantly higher in HC, while patients with severe disease exhibited a lower frequency. Pop6 and Pop11 are CD57-, while Pop7 is CD57+. Related to the CD56^bright^ NK cell subset, both Pop14 and Pop15 were identified. While Pop14 does not express perforin or granzyme B, Pop 15 expresses high levels of these two cytolytic markers. As expected, the frequency of Pop14 was very low in patients with severe disease, while Pop15 was significantly expanded in severe COVID-19.

Despite the classification as innate cells, the discovery of memory properties in NK cells hints the role of this cell type in adaptive immunity and long-term responses (Cerwenka and Lanier, 2016; Lam and Lanier, 2017). In humans adaptive NK cells express NKG2C and lack signaling molecules such as FcRγ (FcεRIγ). It is well known that infection by cytomegalovirus (CMV) induces the expansion of adaptive NK cells (Rölle and Brodin, 2016). At the functional level, they have higher expression of granzyme B, secrete higher levels of IFN-γ and TNF, and are capable of mediating ADCC responses and secrete cytokines against CMV infected cells (Rölle and Brodin, 2016). It has been previously described that in patients with severe COVID-19 there is an increase in the frequency of adaptive NK cells in the circulation (Maucourant et al., 2020). Therefore, we studied this NK cell subset in our cohort of patients. First, we determined the frequencies of NKG2C+, FcRγ- and CD57+NKG2C+ subsets within the CD56^dim^ NK cells (Fig. 7A). Results showed that there were no significant differences in the frequency of these subsets, with the exception of an expansion of NKG2C+ and CD57+NKG2C+ cells from patients with moderate disease. When we simultaneously looked at the expression of the NKG2C and FcRγ markers, we could distinguish four subsets: the NKG2C-FcRγ+ subset that represents the conventional NK cells, and the three different subsets of adaptive NK cells, i.e. NKG2C-FcRγ-, NKG2C+FcRγ- and NKG2C+FcRγ+ (Fig. 7B). The conventional NKG2C-FcRγ+ subset was the largest both in patients and HC. The frequency of these four subsets was similar in patients and HC (Fig. 7B and S10A). Nevertheless, the expression of perforin and granzyme B was higher in each adaptive NK cell subset from COVID-19 patients when compared with HC (Fig. S10B).

**Fig. 7.**
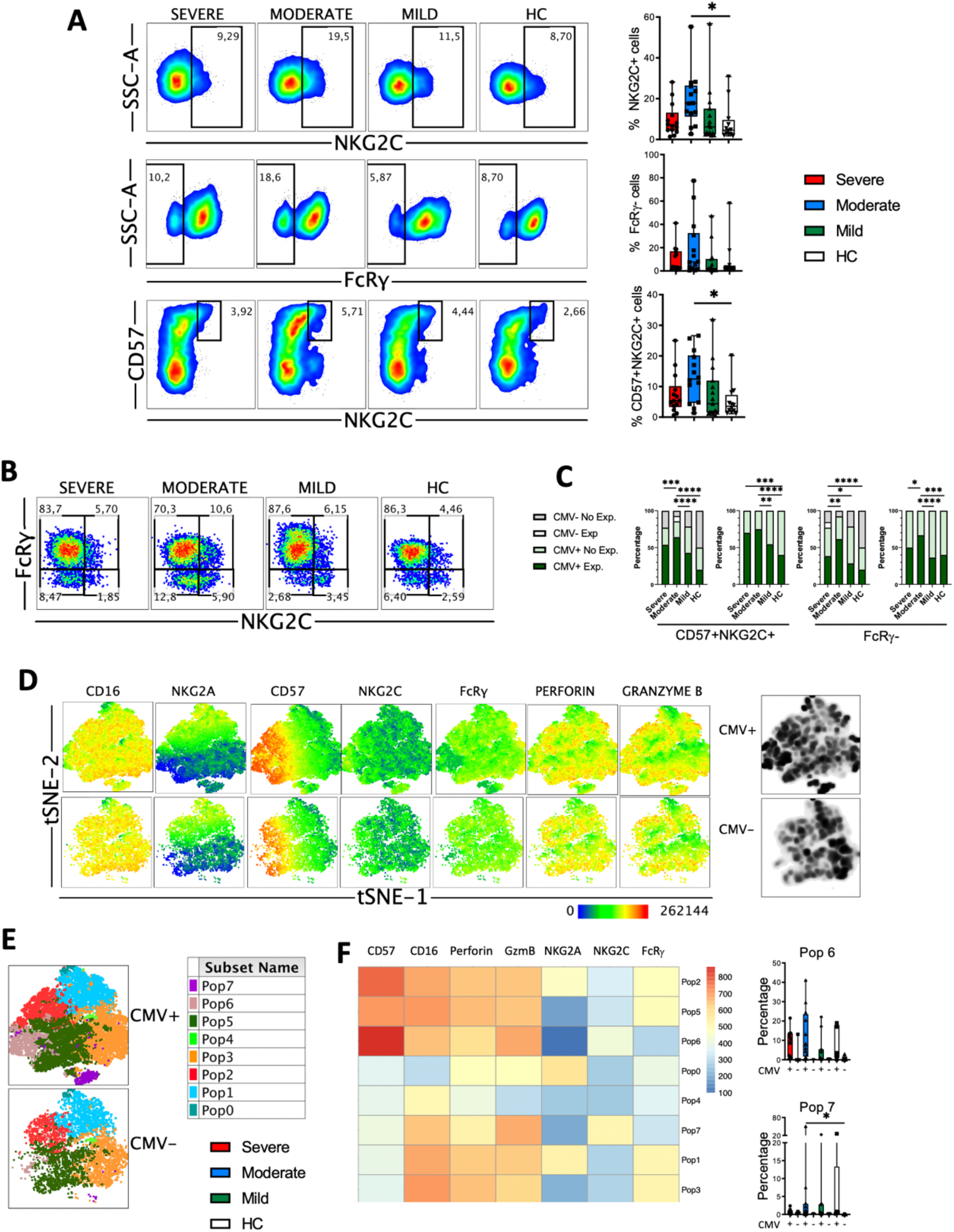
Adaptive NK cells in COVID-19. (**A**) Pseudocolor plots of concatenated CD56^dim^ NK cells from healthy controls (HC) and patients and boxplot graphs of the frequencies of NKG2C+, FcRγ- and CD57+NKG2C+ NK cell subsets. Numbers in the gates are the average of each subset. Boxplots show the median and 25th to 75th percentiles, and the whiskers denote lowest and highest values. Each dot represents a donor. Significance of data was determined by the Kruskal-Wallis test followed by Dunn’s multiple comparison test. *p <0.05. (**B**) Pseudocolor plots of concatenated CD56^dim^ NK cells from healthy controls (HC) and COVID-19 patients showing the expression of NKG2C and FcRγ. Numbers in the quadrants are the average of each subset. (**C**) Percentage of individuals from the indicated groups having or not having adaptive NK cell expansions. Significance of data was determined by chi-squared test. *p <0.05, **p <0.01, ***p <0.001, and ****p <0.0001. (**D**) tSNE projection of the indicated markers and density plots in CD56^dim^ NK cells from CMV-seropositive (CMV+) and CMV-seronegative (CMV-) individuals. (**E**) tSNE projection of CD56^dim^ NK cells populations (Pop) identified by FlowSOM clustering tool from CMV+ and CMV-donors. (**F**) Fluorescence intensity of each Pop as indicated in the column-scaled z-score and boxplot graphs showing the frequencies of Pop6 and Pop7 in CMV+ and CMV-HC and COVID-19 patients. Each dot represents a donor.

It is well known that human adaptive NK cells are expanded in individuals that are infected by CMV (Lam and Lanier, 2017; Rölle and Brodin, 2016). Hence, we studied adaptive NK cells expansion according to the CMV serology status of patients and healthy controls. As reported by Maucourant et al. (Maucourant et al., 2020), we have determined that there is an expansion of adaptive CD57+ NKG2C+ NK cells when they are more than 5% of circulating CD56^dim^ cells. In addition, we have also determined that there is an expansion of adaptive FcRγ-NK cells when they represent more than 7% of the CD56^dim^ cells. Results in Fig. 7C shows that CMV-seronegative individuals do not have expansions of adaptive NK cells, except for one patient with moderate disease, in which the NKG2C+CD57+ cells represented more than 5%, and another patient with severe disease, in which the FcRγ-cells were more than 7%. When only the CMV-seropositive individuals were taken into account, we observed a significant expansion of adaptive NK cells in patients with moderate and severe COVID-19, which was more pronounced when NKG2C+CD57+ cells were taken into account instead of FcRγ-cells (Fig. 7C).

Finally, we performed high-dimensional mapping using tSNE representation and we observed that some regions were preferentially found in CMV-seropositive individuals compared with CMV-seronegative donors (Fig. 7D). Then, to better understand the differences in NK cell subsets between CMV-seropositive and CMV-seronegative individuals we used the FlowSOM clustering tool and compared the expression of seven markers to define 8 populations (Fig. 7E). Using this approach, we were able to identify some populations that were differentially expressed between the two groups of donors (Fig. 7F and S10C). Specifically, the adaptive NK cells Pop6 and Pop7 were characterized by the phenotype NKG2C+FcRγ-, and while Pop6 was CD57+, Pop7 was CD57-. As expected, the frequencies of Pop6 and Pop7 were higher in CMV-seropositive HC and COVID-19 patients, although we did not see significant differences.

### Statistical analysis reveals the relationships between circulating T cells, NK cells and monocytes with disease severity in COVID-19 patients

We first performed a bivariate analysis of 203 clinical laboratory and flow cytometry variables (Table S3). We selected the statistically significant variables for a multivariate analysis. Then, to reduce the number of variables to include in the multivariate analysis we performed a principal component analysis (PCA) (Fig. 8A). Components 1 to 4 explained around 73.7% of the variance, and components 1 and 2 explained around 60.8% of the variance (Fig. S11A). In Figure S11B the contribution of each variable to components 1 to 4 is shown. The expression of CD300 molecules in monocytes and of granzyme B in CD56^bright^ NK cells are variables, along with the frequency of lymphocytes and neutrophils, that significantly contribute to component 1. For component 2, the CD300 receptors expression in monocytes, frequency of neutrophils and lymphocytes, along with CRP, fibrinogen, lymphocyte count and PD-1 in CD4 effector-memory T cells, are variables that contribute significantly. Given that there are three categories (mild, moderate and severe) we performed multinomial logistic regression models. When we compared patients with a mild disease with those with a moderate or severe disease, results showed that component 1 is significantly different between patients with mild and severe disease, while component 2 is significantly different between patients with mild and moderate disease (Fig. 8B, upper panel). On the other hand, when we compared patients with moderate disease with those with a mild and severe disease, we could see that component 1 was significantly different between patients with a moderate and severe disease and, as expected, component 2 was different between patients with moderate and mild disease (Fig. 8B, lower panel).

**Fig. 8.**
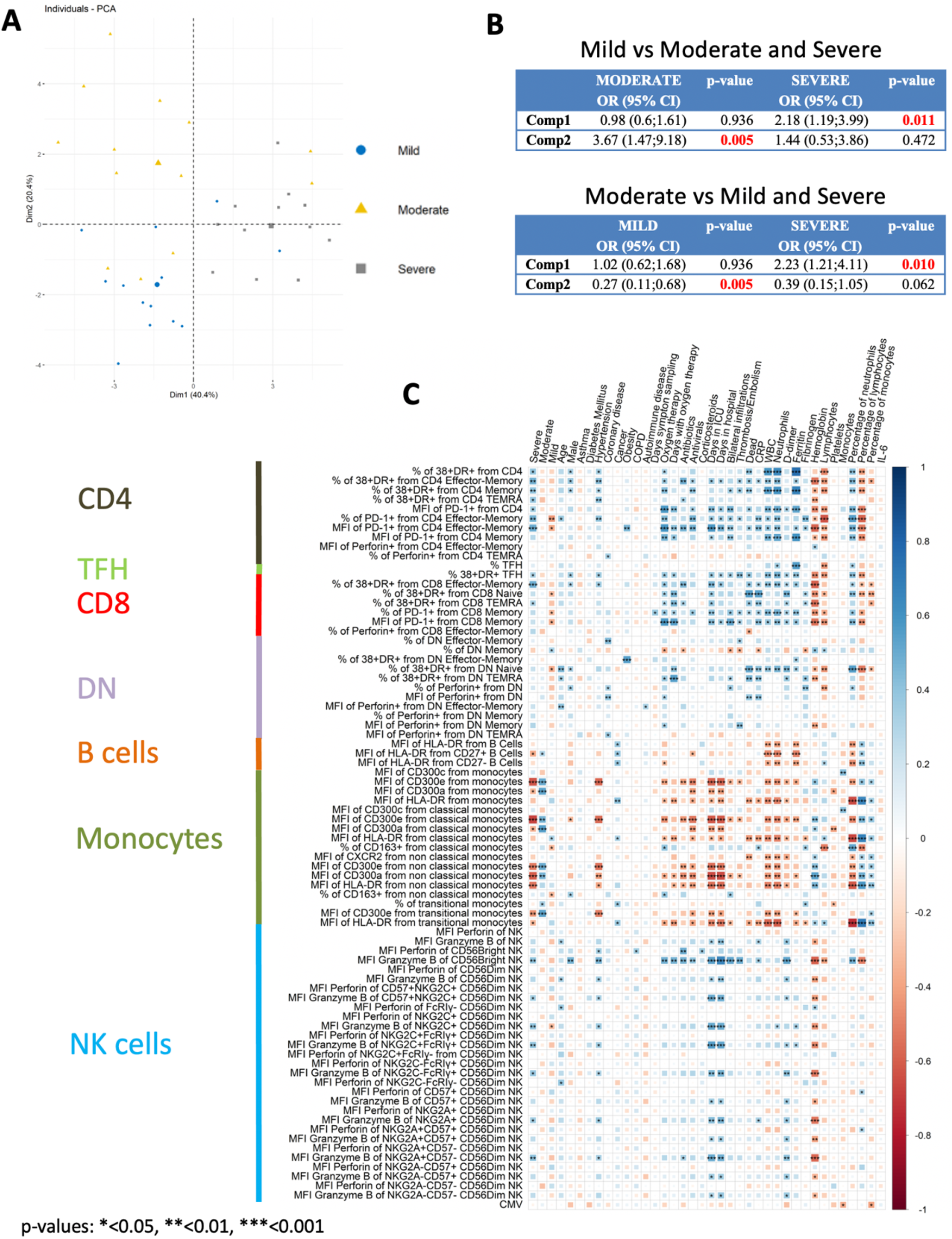
Multivariate analysis and correlation studies of immune cell phenotypes and clinical parameters. (**A**) Representation of the principal component analysis (PCA) results obtained with the most discriminant markers between patients groups. (**B**) Multinomial logistic regression model and statistical significance. Upper: Patients with mild disease versus patients with moderate and severe disease. Lower: Patients with moderate disease versus patients with mild and severe disease. Odd ratio (OR), 95% confidence interval (CI) and p-values are indicated. (**C**) Correlogram showing Spearman correlation of the indicated flow cytometry data and clinical features for COVID-19 patients. Only flow cytometry data that were statistically significant from the bivariate analysis (Table S3) were considered for the analysis.

Next, we performed correlation analysis. A different correlogram pattern was observed between HC and patients groups when we looked at the correlation between the significant flow cytometry variables (Fig. S12). Then, the analysis was performed to look for associations between the general laboratory values, clinical features and degrees of severity with the significant flow cytometry variables (Fig. 8C). Indeed, the correlogram revealed many direct and inverse correlations providing a very valuable insight into the flow cytometry variables that were associated with the degree of severity. Very importantly, a strong direct association of activated T cells and granzyme B expression in CD56^bright^ NK cells with severe disease was observed. Furthermore, and as expected, there was a strong direct association between activated T cells and granzyme B expression in CD56^bright^ NK cells with clinical features of severity such as oxygen therapy, days with oxygen therapy, days in ICU, thrombosis/embolism and bilateral lung infiltrations. Related to clinical laboratory, we observed a direct correlation between activated T cells and granzyme B expression in CD56^bright^ NK cells with ferritin, counts and frequency of neutrophils, D-dimer, and CRP, and an inverse association with hemoglobin and counts and frequency of lymphocytes.

On the other hand, a very interesting picture emerged when the expression of CD300 receptors on monocytes was taken into consideration. A positive correlation between CD300a and CD300e expression with moderate disease was observed. However, a sharp shift in their correlation was detected in patients with severe disease. In fact, an inverse correlation between CD300a and CD300e expression in monocytes with other clinical features of severity was also observed. We also looked at the expression of HLA-DR in monocytes, and there was also an inverse correlation with clinical features associated to severity such as days in ICU, dead, CRP, D-dimer, etc. Altogether, these results indicate that activated T cells, high expression of granzyme B in NK cells and a sharp shift in the expression of CD300 receptors are very significant features of patients with severe COVID-19.

## DISCUSSION

In this study, we have carried out a phenotypic characterization of circulating immune cells in order to determine correlates that may help to distinguish between degrees of severity in COVID-19. In addition, the findings that we have obtained could shed some light into the underlying mechanisms of the disease immunopathogenesis, especially in severe cases. Among other findings, an increased activation status of all T cell subsets (CD4, CD8, circulating TFH and DN cells) and highly armed cytotoxic cells, mostly NK cells and TEMRA and effector-memory T cells, are characteristics of severe disease. Furthermore, to our knowledge, we have uncovered a previously unrecognized alteration and a sharp shift in the pattern of expression of CD300 receptors on monocytes. Very importantly, conventional and unsupervised analyses lead to very similar conclusions highlighting the relevance of our findings.

A possible limitation of our study is the size of the cohort of patients with COVID-19 (n=44) and HC (n=12). However, and to avoid problems related to the diversity of participants, we have been very careful in choosing a cohort that is as uniform as possible before we performed the study. Thus, in this way, it is very important to point out that there are no significant differences between the groups of patients in relation to age, gender or between the number of days since the symptoms onset and samples collection.

It is widely accepted that immune dysregulation contributes to the pathology seen in severe cases of SARS-CoV-2 infection. A consistent finding in many studies, including ours, is that COVID-19 patients display robust activation of the T cell pool, although a considerable portion of patients have minimal levels of activation compared to HC (Mathew et al., 2020). Our study shows that the frequency of activated T cells increases with the degree of severity of the disease. The frequency of activated memory, effectormemory and TEMRA CD4, CD8, DN, and even circulating TFH cells, is increased in patients with severe COVID-19. Furthermore, a very significant correlation was observed between many of these activated T cell subsets with severe disease while, as expected, an inverse association was observed with patients that experienced a mild disease. We also observed that CD4 and CD8 T cells from patients with severe disease tended to have higher levels of perforin. How these activated and perforin containing T cells contribute to the disease pathology is not well known. However, it has been proposed that highly differentiated T cells may induce damage during SARS-CoV-2 infection in a mechanism involving the induction of ligands, as for example MICA/B, for activating NK cell receptors in the respiratory epithelium. The recruited terminally differentiated T cells, which among other things are characterized by the expression of high levels of perforin and NK cell receptors, may recognize and induce T cell receptor-independent killing of epithelial cells in the respiratory tract and lungs (AN and DW, 2020). Therefore, it is tempting to speculate that in patients with severe COVID-19, the expansion of activated T cells, especially the effector-memory and TEMRA cells, have a role in the pathology of the disease. To confirm this affirmation, an in depth analysis of these cell subsets (in blood and tissues) from patients with different degrees of severity is required, including a complete study of their NK cell receptor repertoire. On the other hand, we observed that cytotoxic (perforin containing) CD4 T cells are expanded in a subset of COVID-19 patients. Both protective and pathogenic role for these cells during viral infection have been proposed (Broadley et al., 2017; Sanchez-Martinez et al., 2019). Even more, it has been postulated that an expansion of cytotoxic CD4 T cells drives cardiovascular disease in certain inflammatory conditions and they are triggered by CMV infection (Broadley et al., 2017). There is no doubt that it is necessary to determine if these cytotoxic CD4 T cells have some role in tissue damage and thrombotic complications in COVID-19.

Although we did not find a significant increase in the frequency of circulating TFH, it was possible to observe an expansion in a subset of patients. Nevertheless, circulating TFH were more activated in severe COVID-19, suggesting that these CD38+HLA-DR+ TFH had a recent antigen encounter and may be providing B cell help (Crotty, 2019), possibly as a part of an extrafollicular response, which somehow is explained by the observed low levels of CXCR5 in B cells from patients with severe disease. It has been shown in other viral infections that activated circulating TFH correlates with blood plasmablasts frequency (Crotty, 2019). We did not observe such correlation and, furthermore, we only saw an increase in the plasmablasts frequency in patients with moderate disease. This is in contrast with other studies (Mathew et al., 2020) and we do not know the reason for this discrepancy. Nevertheless, it is important to point out that we have not determined the specific SARS-CoV-2 plasmablast response, but the frequency of total plasmablasts.

Similar to Maucourant et al. (Maucourant et al., 2020), we have also found that COVID-19 patients exhibited expansions of adaptive NK cells as well as highly armed NK cells. Considering that adaptive NK cells are not a uniform NK cell subset (Rölle and Brodin, 2016), it is important to point out that we observed the expansions when we use the two more common gating strategies to identify adaptive NK cells: CD57+NKG2C+ cells and FcRγ-cells. What marker is best for the identification of adaptive NK cells is a matter of discussion, but recently it has been shown that editing the FcRγ gene reprograms conventional NK cells to display functional and phenotypical characteristics of adaptive NK cells (Liu et al., 2020). Expansions of adaptive NK cells occurred mostly, but not exclusively, on those individuals that were CMV-seropositive, raising the question of a possible CMV reactivation in COVID-19. However, the lack of proliferation of CMV specific T cells in SARS-CoV-2 infection and the absence of correlation between the expansions of adaptive NK cells with specific CMV IgG titers argues against the CMV reactivation as the cause (Maucourant et al., 2020; Sekine et al., 2020). Nevertheless, more studies are required to define what drives the expansion of adaptive NK cells in COVID-19 patients.

There was an apparent increase in the levels of perforin and granzyme B in NK cells from COVID-19 patients that correlated with the disease severity. This increased arming of NK cells was very evident in the CD56^bright^ subset, which in patients with severe COVID-19 expressed high levels of perforin and even higher of granzyme B. Interestingly, two populations (Pop14 and Pop15) that differed in the expression of perforin, granzyme B and CD16 were identified within the CD56^bright^ NK cells. Some authors have previously suggested that CD56^bright^CD16+ NK cells represent a more mature stage within the CD56^bright^ subset (Campos et al., 2015). Therefore, we could conclude that in severe COVID-19 there is a shift toward a more mature NK cell within the CD56^bright^ cells. On the other hand, we did not observe a difference in the maturation status of the CD56^dim^ subset according to the expression of NKG2A and CD57.

Correlation studies showed a strong association of granzyme B expression in CD56^bright^ NK cells with severe disease as shown by clinical features such as bilateral infiltrations, thrombosis/embolism, oxygen therapy, neutrophilia, etc. A less strong association was also observed when the granzyme B expression in CD56^dim^ NK cells was taken into account. The presence of these highly armed NK cells in patients with severe disease may suggest the possibility to eliminate more efficiently target cells, including virus-infected cells, and activated T cells that may cause immunopathology, and therefore modulate the adaptive immune response (Waggoner et al., 2012). But also, these NK cells can cause tissue damage in a way similar to how respiratory syncytial virus causes acute lung damage (Li et al., 2012). Undoubtedly, more studies are required to know how these two aspects of NK cells contribute to the immunopathogenesis of COVID-19. A decrease in non-classical monocytes and an increase in the transitional subset have been associated to a severe and mild COVID-19, respectively (Sánchez-Cerrillo et al., 2020; Schulte-Schrepping et al., 2020; Silvin et al., 2020). Although we did not find significant differences between patients and HC in the frequency of monocyte subsets, a tendency to a diminution in the frequency of the non-classical subset and an increase in the transitional monocytes was evident. In agreement with previous results (Giamarellos-Bourboulis et al., 2020; Schulte-Schrepping et al., 2020; Silvin et al., 2020), we observed a decrease in HLA-DR expression in monocytes. The decrease was gradual and it was very significant in patients with severe COVID-19. HLA-DR^low^ monocytes are considered dysfunctional and are an established surrogate marker of immunosuppression in sepsis (Venet et al., 2020). Acute infections trigger an emergency myelopoiesis that is characterized by the mobilization of immature myeloid cells, which are linked to immunosuppressive functions (Loftus et al., 2018). Therefore, it is reasonable to propose that an increment in dysfunctional monocytes and an emergency myelopoiesis are factors that contribute to the development of severe disease.

The CD300 molecules are type I transmembrane proteins expressed on the surface of immune cells and are divided in two groups: activating and inhibitory receptors (Borrego, 2013). CD300 receptors are able to bind different ligands, mostly lipids such as phosphatidylserine, phosphatidylethanolamine and ceramide (Borrego, 2013; Izawa et al., 2014; Simhadri et al., 2012; Zenarruzabeitia et al., 2015). They regulate many signaling pathways, as for example monocyte and neutrophil activation (Alvarez et al., 2008; Borrego, 2013; Simhadri et al., 2013; Zenarruzabeitia et al., 2015, 2016). The importance of CD300 molecules in several pathological conditions has been highlighted by multiple studies describing the role of this family of receptors in allergic disorders, autoimmune and inflammatory diseases, cancer, sepsis and viral infections (Borrego, 2013; Vitallé et al., 2019; Zenarruzabeitia et al., 2015). In this study we have observed a very intriguing expression pattern of the CD300 molecules in COVID-19 patients. The expression of CD300a and CD300e gradually increased in monocytes and granulocytes from patients with mild and moderate disease. It is well known that activation of neutrophils and monocytes with pro-inflammatory stimuli, such as LPS and GM-CSF, increased the expression of CD300a and CD300e (Alvarez et al., 2008; Zenarruzabeitia et al., 2016). Therefore, it is plausible to assume that in patients with moderate and mild disease the increased expression in these two CD300 receptors is a consequence of the inflammatory milieu. However, a sudden change in the pattern of CD300a and CD300e receptors expression happened in patients with severe COVID-19 that exhibited very low levels of these two molecules. The cause for this sharp shift is not known. However, some clues could be found in the HL-60 acute myeloid leukemia cell model to study the differentiation towards monocytes and neutrophils (Alvarez et al., 2008). Undifferentiated HL-60 cell do not express CD300 molecules, but when neutrophil and monocyte differentiation was induced, a significant increase in CD300a cell surface expression was observed, indicating that the CD300a receptor expression is developmentally regulated (Alvarez et al., 2008). Therefore, a possible explanation for the sharp decrease in CD300a expression, and also possibly CD300e, is that the circulating monocytes and granulocytes are more immature (and possibly more dysfunctional) that those observed in mild and moderate COVID-19. This is somehow supported by the lower HLA-DR expression in monocytes from patients with severe disease. Evidently, more studies are required to understand not only the exact mechanisms involved in the regulation of CD300 receptors expression in COVID-19, but also the role that they play in this disease. Besides that, the determination of CD300 receptors expression in COVID-19 is very important as the statistical, including PCA analysis, and correlation studies show.

In conclusion, the two important findings of this study were that the unsupervised analysis obtained similar results and reached similar conclusions than the conventional analysis, and that the statistical and correlation studies corroborated that several immune alterations were in close relationship with the severity of the disease as shown by the clinical features and clinical laboratory data. This study could be improved in the future by increasing the number of recruited patients, performing longitudinal studies, analyzing in more depth immune cell subsets, as for example terminal differentiated T cells and monocytes, from blood and tissues, obtaining more comprehensive clinical data, etc. Still, our study, along with those published by others, provides a compilation of immune response data that, at least, could help in two ways: first, by providing additional light on the immune mechanisms behind the development of severe COVID-19, and second, with the potential benefit of helping clinicians to decide which therapeutic approach is better for each patient.

## PATIENTS, MATERIALS AND METHODS

### Patients and healthy donors

In this study, we used plasma and whole blood samples cryopreserved in Cytodelics Stabiliser (http://www.cytodelics.com) from SARS-CoV-2 infected patients (n=44) and adult healthy donors (n=12). Patients were recruited at Cruces University Hospital. All samples were collected through the Basque Biobank for Research (https://www.biobancovasco.org). The Basque Biobank complies with the quality management, traceability and biosecurity, set out in the Spanish Law 14/2007 of Biomedical Research and in the Royal Decree 1716/2011. Patients recruited into the study tested PCR-positive for SARS-CoV-2, except one patient with mild disease, from March to June 2020 and were classified in 3 different clinical severity groups: mild, moderate and severe, according to clinical criteria (see Table S1 and S2). All donors provided written and signed informed consent in accordance with the Declaration of Helsinki. This study was approved by the Basque Ethics Committee for Research with Medicines (CEIm-E) with the number CES-BIOEF 2020-13.

### Cell preparation and flow cytometry

From each donor a whole blood sample was collected in EDTA, from which one aliquot was centrifuged to obtain plasma and another aliquot was frozen using Cytodelics whole blood cell stabilizer. For extracellular staining, 200μL of blood + stabilizer mixture thawed at 37°C were incubated with the specific antibodies for 23 min at room temperature (RT) in the dark. Next, red blood cells were lysed using 2mL of 1X BD FACS Lysing Solution (BD Biosciences) for 12 min at RT. Cells were washed with PBS containing 2.5% of bovine serum albumin (BSA) (Sigma-Aldrich) and were permeabilized and fixed using Cytofix/Cytoperm Plus Kit (BD Biosciences) following manufacturer’s instructions. Then, intracellular staining was performed using the corresponding antibodies during 30 min at 4°C in the dark. Lastly, cells were washed, resuspended in 250 μL of PBS and acquired in a LSRFortessa X-20 flow cytometer (BD Biosciences). Three flow cytometry panels were used to study T and B cells, NK cells and monocytes (see Table S4).

### Flow cytometry data analysis

FCS 3.0 files were exported from the FACSDiva and imported into FlowJo v.10.7.1. for subsequent analysis. The following plug-ins were used: DownSample (1.1), tSNE and FlowSOM (2.6). Manual and automated analyses were performed. For the automated analysis, events were first downsampled from the gates of interest (CD4 T cells, CD8 T cells, B cells, monocytes, NK cells and CD56^dim^ NK cells) across all samples using DownSample plug-in. Then, downsampled populations were concatenated for the analysis. tSNE was run using the parameters indicated in each figure and represented as heatmap and density plot. FlowSOM was run using the same parameters from the tSNE panels.

### Clinical laboratory data

Hemograms and serum determinations (D-dimer, ferritin, fibrinogen, CRP, etc.) from all patients were realized at the clinical laboratory of Cruces University Hospital. In addition, IL-6 levels and CMV serology were determined in plasma samples, also at the clinical laboratory of Cruces University Hospital. Samples for all determinations were obtained the same day than samples for flow cytometry analysis.

### Statistical analysis and data representation

GraphPad Prism v.8.4.3 was used for graphical representation and statistical analysis. Data were represented as boxplots with the median and 25^th^ to 75^th^ percentiles, and the whiskers denote lowest and highest values. Each dot represented a donor. Significance was determined by the Kruskal-Wallis test adjusting for multiple comparisons using Dunn’s test. For categorical comparisons, the significance was determined by chi-squared test.

Correlation plots between variables were calculated and visualized as correlograms using R function *corrplot.* Spearman’s Rank Correlation coefficient was indicated by square size and heat scale. Significance was indicated by *p <0.05, **p <0.01, ***p <0.001, and ****p <0.0001. p values were adjusted using the Benjamini & Hochberg test.

Bivariate analyses were performed (Table S3). First, using the Shapiro-Wilks normality test we determined if variables followed a normal distribution. If they did, the average and standard deviation were reported, and if they did not have a normal distribution the median and interquartile range were indicated. To determine statistical significance, we use the Student’s t test if the variable follows a normal distribution and the Mann-Whitney U test otherwise. To take into account multiple comparisons, we also presented the adjusted p-values using the Benjamini & Hochberg test (Table S3). Multivariate analysis was performed after selecting the statistically significant variables from bivariate analysis. To reduce the number of variables to include in the multivariate analysis we performed a Principal Component Analysis (PCA). Multinomial logistic regression models were performed to determine the components that were associated with the disease.

## Acknowledgements

We thank all patients and healthy controls who participated in this study and the staff from the Basque Biobank for Research. This work was supported by a grant from the “Agencia Estatal de Investigation” Project PID2019-109583RB-I00/AEI/10.13039/501100011033. OZ is recipient of a postdoctoral contract funded by “Instituto de Salud Carlos III-Contratos Sara Borrell 2017 (CD17/0128)” and the European Social Fund (ESF)-The ESF invests in your future. GA-P is recipient of a fellowship from the BBK Fundazioa (1543/2006_0001) and from the Jesús de Gangoiti Barrera Foundation (FJGB20/002). IT is recipient of a predoctoral contract funded by the Department of Education, Basque Government (PRE_2019_2_0109). AO is recipient of a fellowship from the Jesús de Gangoiti Barrera Foundation (FJGB19/002). FB is an Ikerbasque Research Professor, Ikerbasque, Basque Foundation for Science.

## Author contribution

FB conceived the project; OZ and FB designed experiments; IS-B, JN-A, NI-A, and EA-A obtained the clinical samples and clinical data from COVID-19 patients; OZ and FB obtained samples from healthy controls; RP-G determined IL-6 and CMV serology from patients and healthy controls; OZ stained and acquired flow cytometry samples; GA-P, and FB performed flow cytometry analysis; GA-P, and SP-F performed computational and statistical analysis; GA-P, SP-F and FB compiled figures; OZ, GA-P, IT, and AO provided intellectual input; FB wrote the manuscript; all authors reviewed the manuscript.

