## Supplementary Material for "T cell activation, highly armed cytotoxic cells and a sharp shift in monocytes CD300 receptors expression is characteristic of patients with severe COVID-19"

**This file includes:**

Fig. S1. Gating strategy to identify immune cells.
Fig. S2. T cell subsets and CD4/CD8 ratio in COVID-19.
Fig. S3. CD4 T cells activation status, PD-1 and perforin expression and CD4 T follicular helper (TFH) cells in COVID-19 patients.
Fig. S4. Expression of CXCR5 and HLA-DR in B cell subsets.
Fig. S5. Activation status, PD-1 and perforin expression in CD8 T cells from COVID-19 patients.
Fig. S6. Activation status and perforin expressing double negative (DN) T cells in COVID-19 patients.
Fig. S7. Expression of CD300a in CD66b+ cells (granulocytes).
Fig. S8. Monocyte populations identified by FlowSOM clustering.
Fig. S9. Perforin and granzyme B expression on NK cell subsets and populations identified by FlowSOM clustering.
Fig. S10. Adaptive NK cells: subsets, perforin and granzyme B expression and populations identified by FlowSOM clustering.
Fig. S11. Components and variables for the PCA.
Fig. S12. Correlation between significant flow cytometry variables.
Table S1. Clinical characteristics of COVID-19 patients.
Table S2. Clinical laboratory results of COVID-19 patients.
Table S3. Flow cytometry panels.
Table S4. Bivariate analysis of clinical laboratory and flow cytometry data.

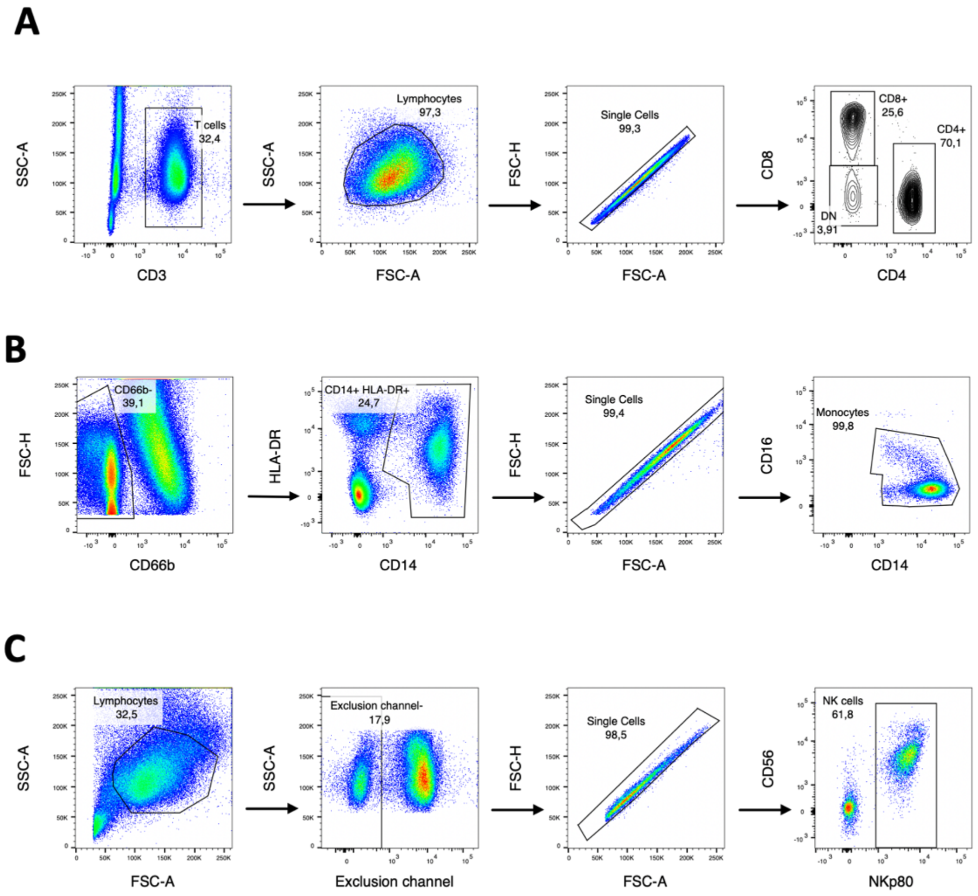

**Fig. S1. Gating strategy to identify immune cells.** Pseudocolor and contour plots representing the gating strategy utilized for the identification of T cells (**A**), monocytes (**B**) and NK cells (**C**). The Exclusion channel included BV510 conjugated antibodies against CD3, CD14 and CD19.

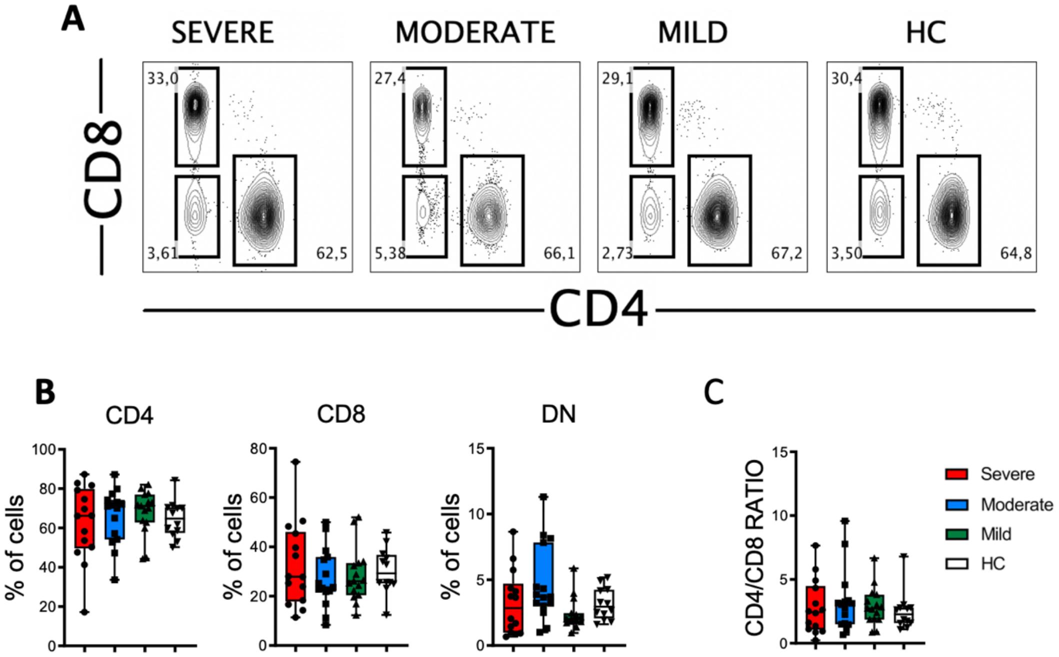

**Fig. S2. T cell subsets and CD4/CD8 ratio in COVID-19.** (**A**) Contour plots of concatenated CD4, CD8 and double negative (DN) T cell subsets from healthy controls (HC) and patients with mild, moderate and severe COVID-19. Numbers in the gates are the average of each subset. (**B**) Boxplot graphs of CD4, CD8 and DN T cell subsets. (**C**) Boxplot graph of CD4/CD8 ratio. Boxplots show the median and 25$^{th}$ to 75$^{th}$ percentiles, and the whiskers denote lowest and highest values. Each dot represents a donor.

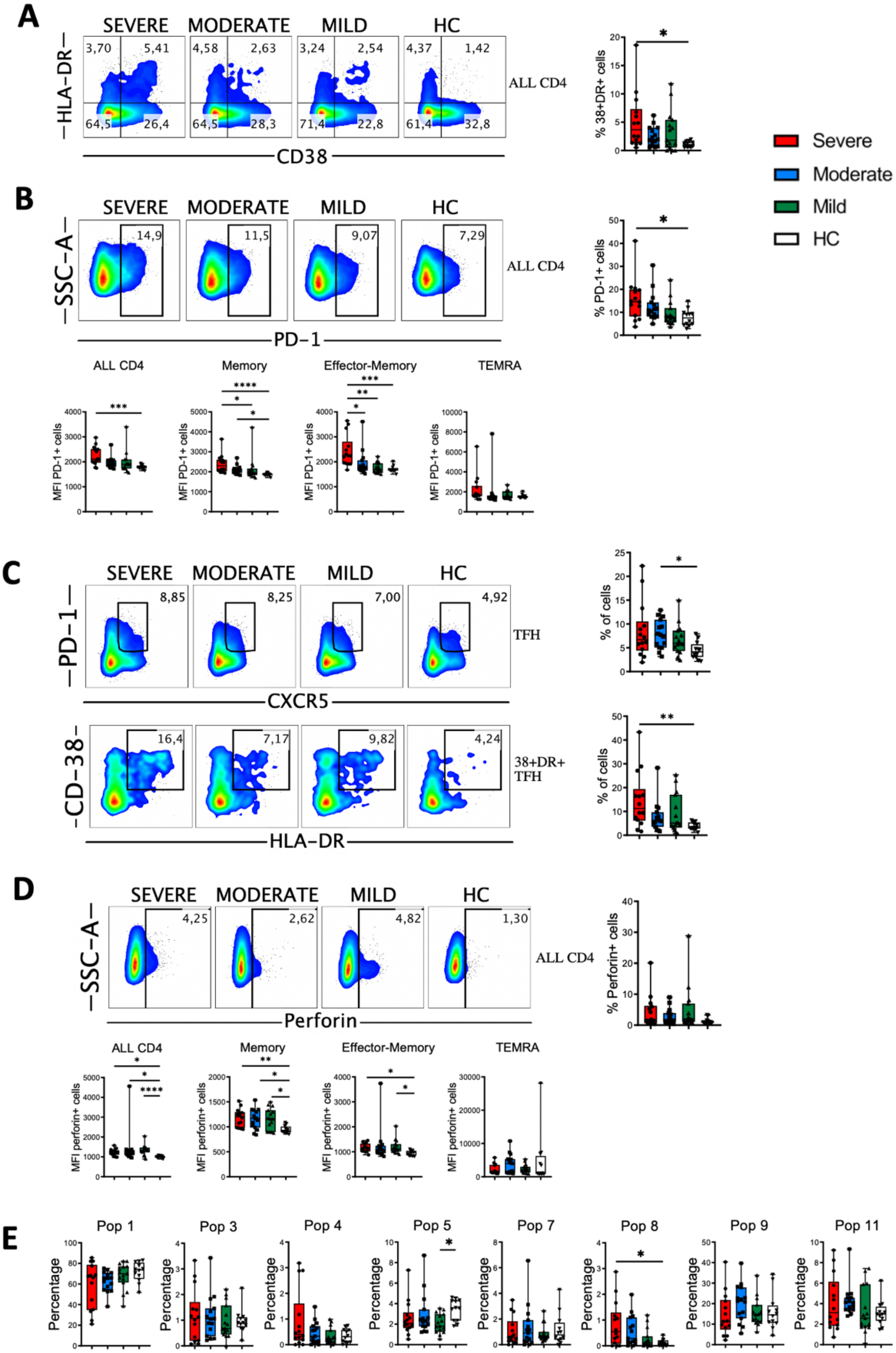
A
SEVERE
MODERATE
MILD
HC
3,70
5,41
4,58
2,63
3,24
2,54
4,37
1,42
64,5
26,4
64,5
28,3
71,4
22,8
61,4
32,8
HLA-DR
CD38
ALL CD4
% 38+DR+ cells
Severe
Moderate
Mild
HC
B
SEVERE
MODERATE
MILD
HC
14,9
11,5
9,07
7,29
SSC-A
PD-1
ALL CD4
% PD-1+ cells
ALL CD4
Memory
Effector-Memory
TEMRA
MFI PD-1+ cells
C
SEVERE
MODERATE
MILD
HC
8,85
8,25
7,00
4,92
PD-1
CXCR5
TFH
% of cells
16,4
7,17
9,82
4,24
CD-38
HLA-DR
38+DR+ TFH
D
SEVERE
MODERATE
MILD
HC
4,25
2,62
4,82
1,30
SSC-A
Perforin
ALL CD4
% Perforin+ cells
ALL CD4
Memory
Effector-Memory
TEMRA
MFI perforin+ cells
E
Pop 1
Pop 3
Pop 4
Pop 5
Pop 7
Pop 8
Pop 9
Pop 11
Percentage

**Fig. S3. CD4 T cells activation status, PD-1 and perforin expression and CD4 T follicular helper (TFH) cells in COVID-19 patients.** (**A**) Pseudocolor plots of concatenated peripheral CD4 T cells from healthy controls (HC) and patients and boxplot graph showing the frequency of all activated cells, which are identified by the coexpression of CD38 and HLA-DR. Numbers in the quadrants are the average of each subset. (**B**). Upper part: pseudocolor plots of concatenated CD4 T cells and boxplot graph of the frequency of all CD4 T cells that are PD-1+. Numbers in the gates are the average of PD-1+ cells in each group. Lower part: boxplot graphs showing the median fluorescence intensity (MFI) of PD-1+ cells in all and CD4 T cell subsets. (**C**) Upper part: pseudocolor plots of concatenated CD4 T cells and boxplot graph representing the frequency of T follicular helper (TFH) cells in HC and COVID-19 patients. Numbers in the gates are the average of TFH cells. Lower part: pseudocolor plots of concatenated TFH CD4 T cells and boxplot graph of activated (CD38+HLA-DR+) TFH cells in HC and patients. Numbers in the gates are the average of activated TFH cells. (**D**) Upper part: pseudocolor plots of concatenated CD4 T cells and boxplot graph showing the frequency of all CD4 T cells that are perforin positive. Numbers in the gates are the average of perforin positive cells in each group. Lower part: boxplot graphs showing the MFI of perforin positive cells in all and CD4 T cell subsets. (**E**) Boxplot graphs representing the frequencies of Pop1, Pop3, Pop4, Pop5, Pop7, Pop8, Pop9 and Pop11 in HC and COVID-19 patients. Boxplots show the median and $25^{th}$ to $75^{th}$ percentiles, and the whiskers denote lowest and highest values. Each dot represents a donor. Significance of data was determined by the Kruskal-Wallis test followed by Dunn´s multiple comparison test. *p <0.05, **p <0.01, ***p <0.001, and ****p <0.0001.

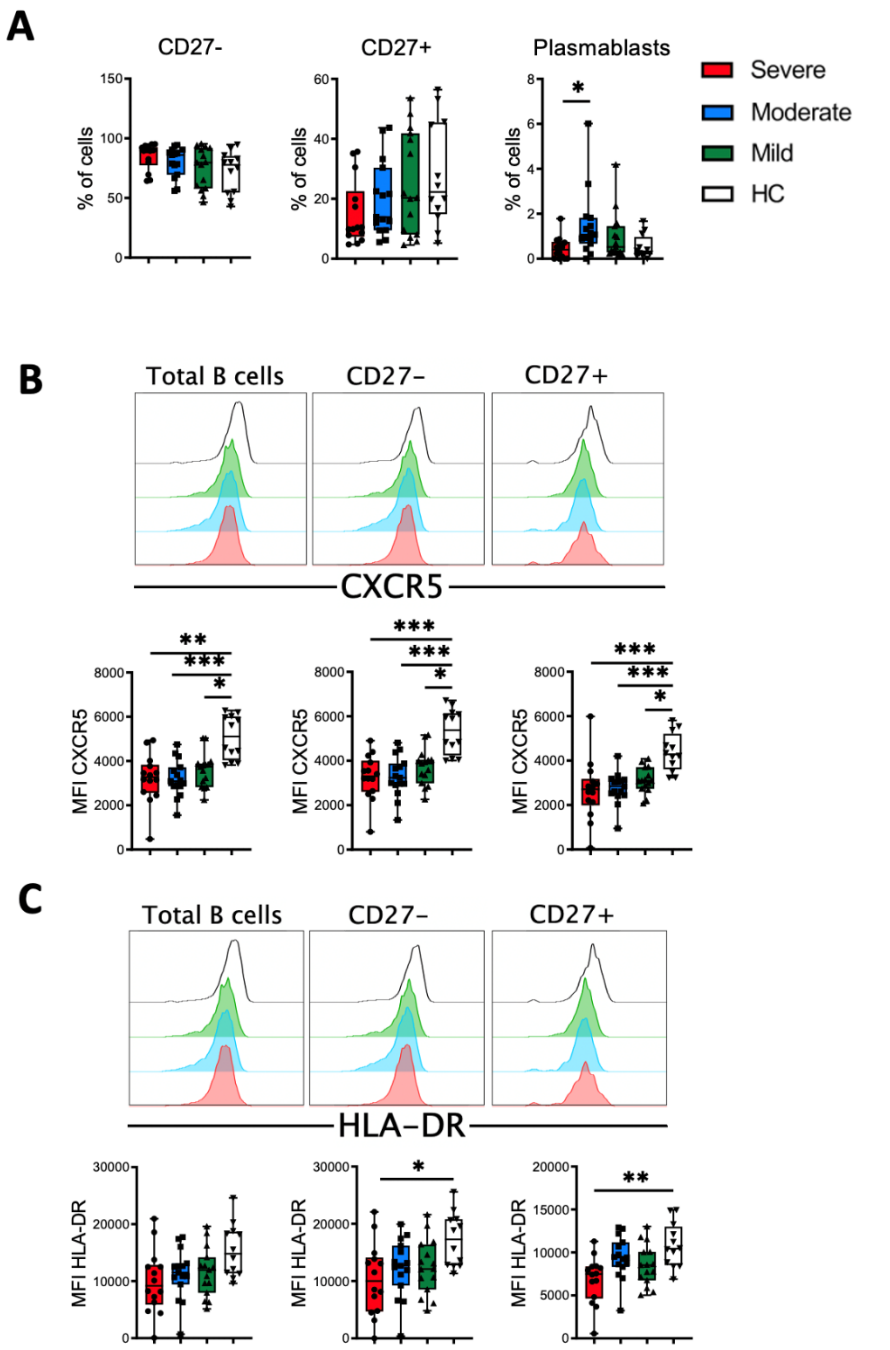

**Fig. S4. Expression of CXCR5 and HLA-DR in B cell subsets.** (**A**) Boxplot graphs of the frequencies of CD27-, CD27+ and plasmablasts (CD27+CD38+) B cell subsets in healthy controls (HC) and COVID-19 patients. (**B**) Histograms of concatenated total B cells, CD27- and CD27+ subsets (upper) and boxplot graphs (lower) representing the median fluorescence (MFI) of CXCR5 in each cell subset. (**C**) Histograms of concatenated total B cells, CD27- and CD27+ subsets (upper) and boxplot graphs (lower) showing the MFI of HLA-DR in each cell subset. Boxplots show the median and 25$^{th}$ to 75$^{th}$ percentiles, and the whiskers denote lowest and highest values. Each dot represents a donor. Significance of data was determined by the Kruskal-Wallis test followed by Dunn´s multiple comparison test. *p <0.05, **p <0.01, ***p <0.001, and ****p <0.0001.

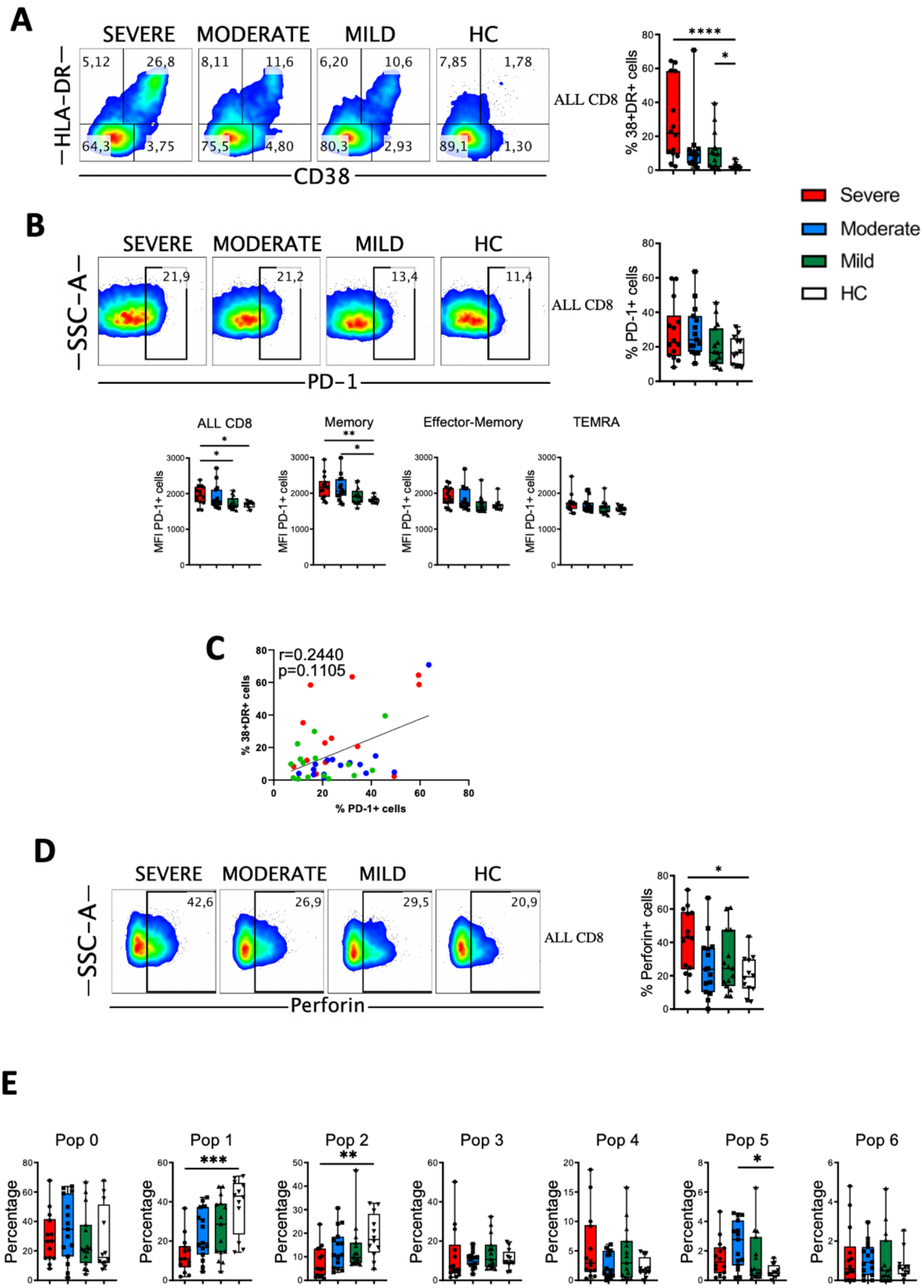
A
SEVERE
MODERATE
MILD
HC
HLA-DR
CD38
5,12
26,8
64,3
3,75
8,11
11,6
75,5
4,80
6,20
10,6
80,3
2,93
7,85
1,78
89,1
1,30
ALL CD8
% 38+DR+ cells
****
*
Severe
Moderate
Mild
HC
B
SSC-A
PD-1
21,9
21,2
13,4
11,4
% PD-1+ cells
ALL CD8
Memory
Effector-Memory
TEMRA
MFI PD-1+ cells
C
r=0.2440
p=0.1105
% 38+DR+ cells
% PD-1+ cells
D
Perforin
42,6
26,9
29,5
20,9
% Perforin+ cells
E
Pop 0
Pop 1
Pop 2
Pop 3
Pop 4
Pop 5
Pop 6
Percentage
***
**

**Fig. S5. Activation status, PD-1 and perforin expression in CD8 T cells from COVID-19 patients.** (**A**) Pseudocolor plots of concatenated peripheral CD8 T cells from healthy controls (HC) and patients and boxplot graph showing the frequency of all activated cells, which are identified by the coexpression of CD38 and HLA-DR. Numbers in the quadrants are the average of each subset. (**B**). Upper part: pseudocolor plots of concatenated CD8 T cells and boxplot graph of the frequency of all CD8 T cells that are PD-1+. Numbers in the gates are the average of PD-1+ cells in each group. Lower part: boxplot graphs showing the median fluorescence intensity (MFI) of PD-1+ cells in all and CD8 T cell subsets. (**C**) Spearman correlation of activated (CD38+HLA-DR+) with PD-1+ CD8 T cells from patients with mild, moderate and severe COVID-19. (**D**) Pseudocolor plots of concatenated CD8 T cells and boxplot graph showing the frequency of all CD8 T cells that are perforin positive. Numbers in the gates are the average of perforin positive cells in each group. (**E**) Boxblot graphs representing the frequencies of Pop0, Pop1, Pop2, Pop3, Pop4, Pop5 and Pop6 in HC and COVID-19 patients. Boxplots show the median and $25^{th}$ to $75^{th}$ percentiles, and the whiskers denote lowest and highest values. Each dot represents a donor. Significance of data was determined by the Kruskal-Wallis test followed by Dunn´s multiple comparison test. *p <0.05, **p <0.001, ***p <0.001, and ****p <0.0001.

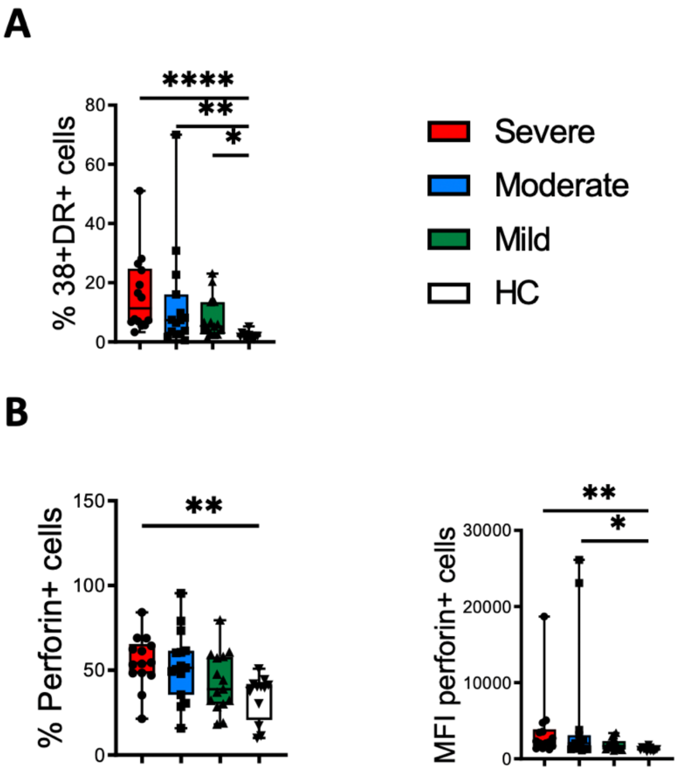

**Fig. S6. Activation status and perforin expressing double negative (DN) T cells in COVID-19 patients.** (**A**) Boxplot graph of the frequency of activated DN T cells in healthy controls (HC) and patients. Activated cells are identified by the coexpression of CD38 and HLA-DR. (**B**) Boxplot graphs representing perforin positive (left) and median fluorescence intensity (MFI) of perforin positive cells (right) in DN T cells. Boxplots show the median and 25$^{th}$ to 75$^{th}$ percentiles, and the whiskers denote lowest and highest values. Each dot represents a donor. Significance of data was determined by the Kruskal-Wallis test followed by Dunn´s multiple comparison test. *p <0.05, **p <0.01, and ****p <0.0001.

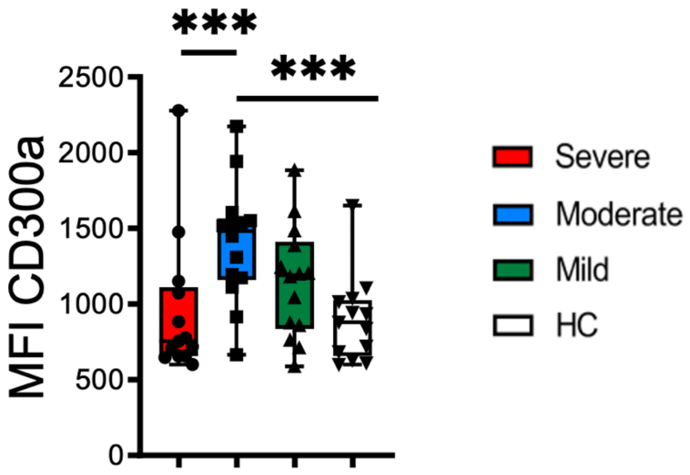

**Fig. S7. Expression of CD300a in CD66b+ cells (granulocytes).** Boxplot graph of the median fluorescence intensity (MFI) of CD300a in CD66b+ cells from healthy controls (HC) and COVID-19 patients. Boxplots show the median and 25$^{th}$ to 75$^{th}$ percentiles, and the whiskers denote lowest and highest values. Each dot represents a donor. Significance of data was determined by the Kruskal-Wallis test followed by Dunn´s multiple comparison test. ***p <0.001.

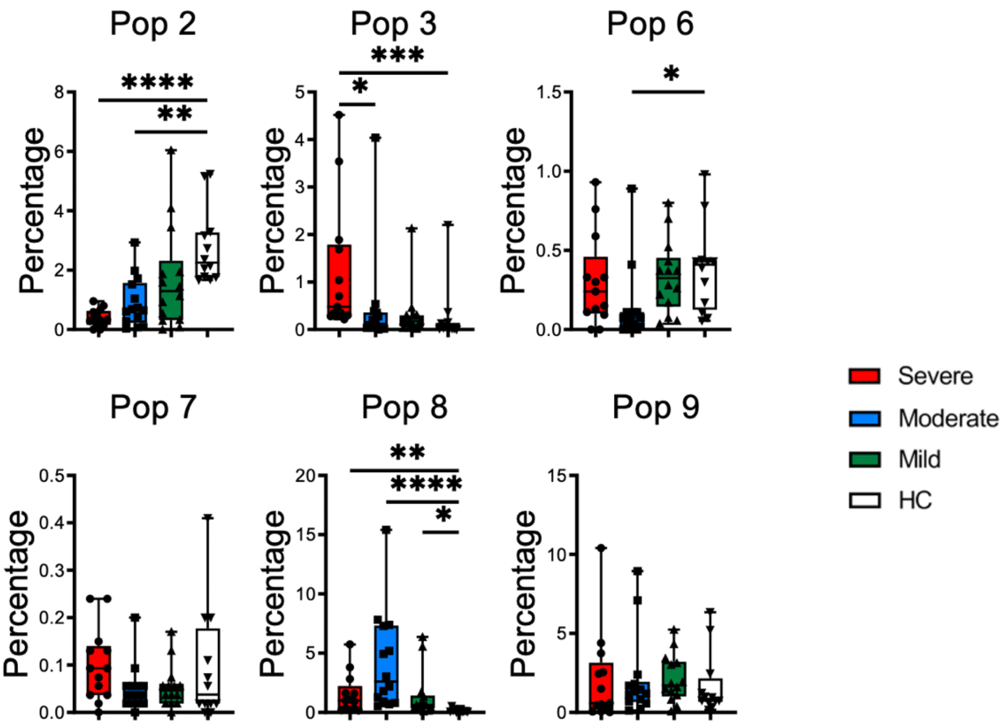

**Fig. S8. Monocyte populations identified by FlowSOM clustering.** Boxplot graphs representing the frequencies of monocytes Pop2, Pop3, Pop6, Pop7, Pop8 and Pop9 from healthy controls (HC) and COVID-19 patients. Boxplots show the median and 25$^{th}$ to 75$^{th}$ percentiles, and the whiskers denote lowest and highest values. Each dot represents a donor. Significance of data was determined by the Kruskal-Wallis test followed by Dunn´s multiple comparison test. *p <0.05, **p <0.001, ***p <0.001, and ****p <0.0001.

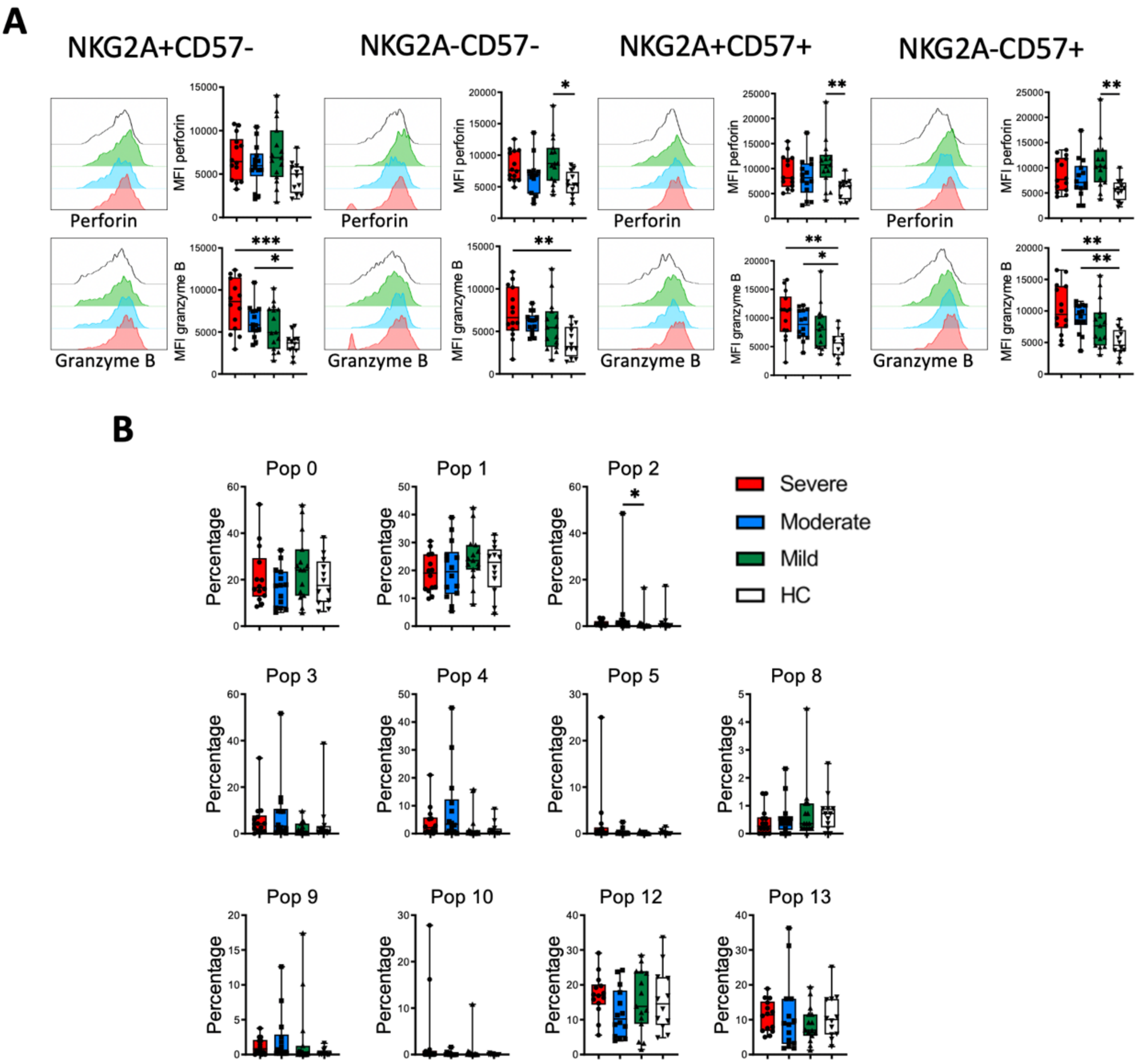

**Fig. S9. Perforin and granzyme B expression in NK cell subsets and populations identified by FlowSOM clustering.** (**A**) Histograms of concatenated CD56dim NK cells and boxplot graphs showing the median fluorescence intensity (MFI) of perforin and granzyme B in each NK cell subset. (**B**) Boxplot graphs representing the frequencies of NK cells Pop2, Pop3, Pop6, Pop7, Pop8 and Pop9 from healthy controls (HC) and COVID-19 patients. Boxplots show the median and 25th to 75th percentiles, and the whiskers denote lowest and highest values. Each dot represents a donor. Significance of data was determined by the Kruskal-Wallis test followed by Dunn´s multiple comparison test. *p <0.05, **p <0.01, and ***p <0.001.

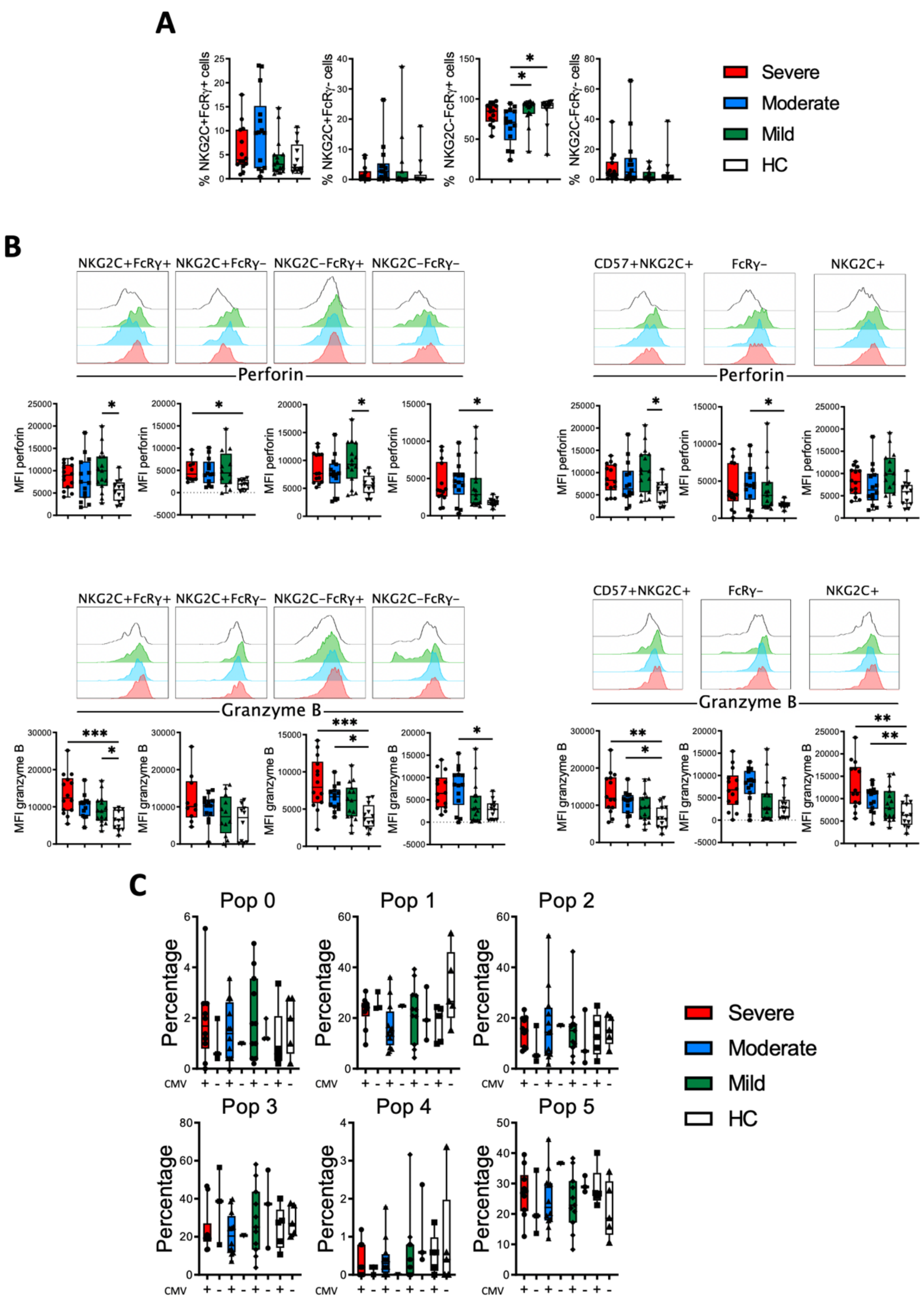

A
% NKG2C+FcRγ+ cells
% NKG2C+FcRγ- cells
% NKG2C-FcRγ+ cells
% NKG2C-FcRγ- cells
Severe
Moderate
Mild
HC
B
NKG2C+FcRγ+
NKG2C+FcRγ−
NKG2C−FcRγ+
NKG2C−FcRγ−
CD57+NKG2C+
FcRγ−
NKG2C+
Perforin
MFI perforin
Granzyme B
MFI granzyme B
C
Pop 0
Pop 1
Pop 2
Pop 3
Pop 4
Pop 5
Percentage
CMV

**Fig. S10. Adaptive NK cells: subsets, perforin and granzyme B expression and populations identified by FlowSOM clustering.** (**A**) Boxplot graphs showing the frequency of NKG2+CFcRγ+, NKG2C+FcRγ-, NKG2C-FcRγ+ and NKG2-CFcRγ- subsets within the CD56dim NK cells in healthy controls (HC) and COVID-19 patients. (**B**) Histograms of concatenated CD56dim NK cells and boxplot graphs showing the median fluorescence intensity (MFI) of perforin (upper panel) and granzyme B (lower panel) in each NK cell subset. (**C**) Boxplot graphs representing the frequencies of NK cells Pop0, Pop1, Pop2, Pop3, Pop4 and Pop5 from CMV-seropositive and CMV-seronegative healthy controls (HC) and COVID-19 patients. Boxplots show the median and 25th to 75th percentiles, and the whiskers denote lowest and highest values. Each dot represents a donor. Significance of data was determined by the Kruskal-Wallis test followed by Dunn´s multiple comparison test. *p <0.05, **p <0.01, and ****p <0.0001.

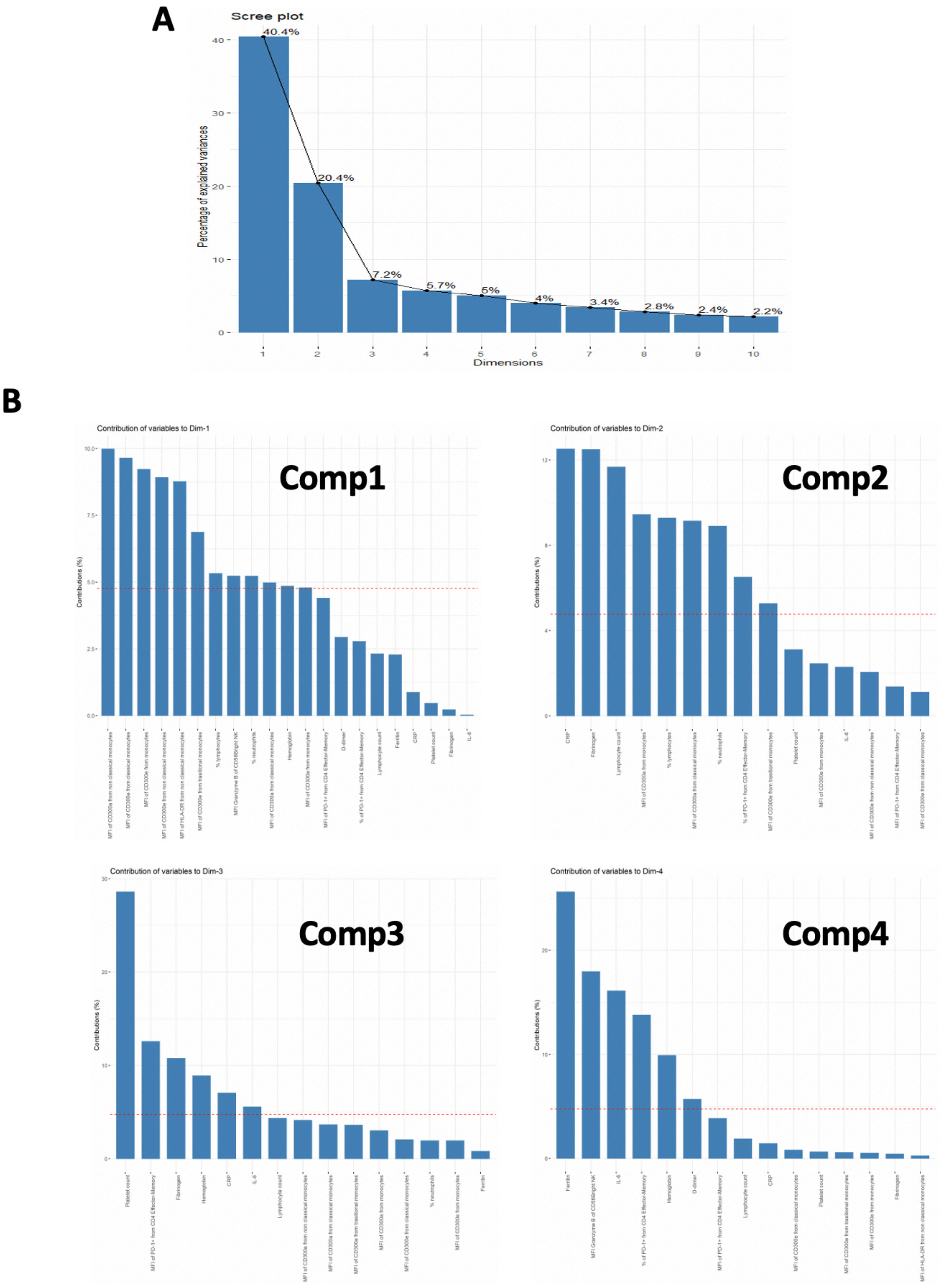

**Fig. S11. Components and variables for the PCA.** (**A**) Components that explain the variance. (**B**) Contribution of the variables to components 1 to 4.

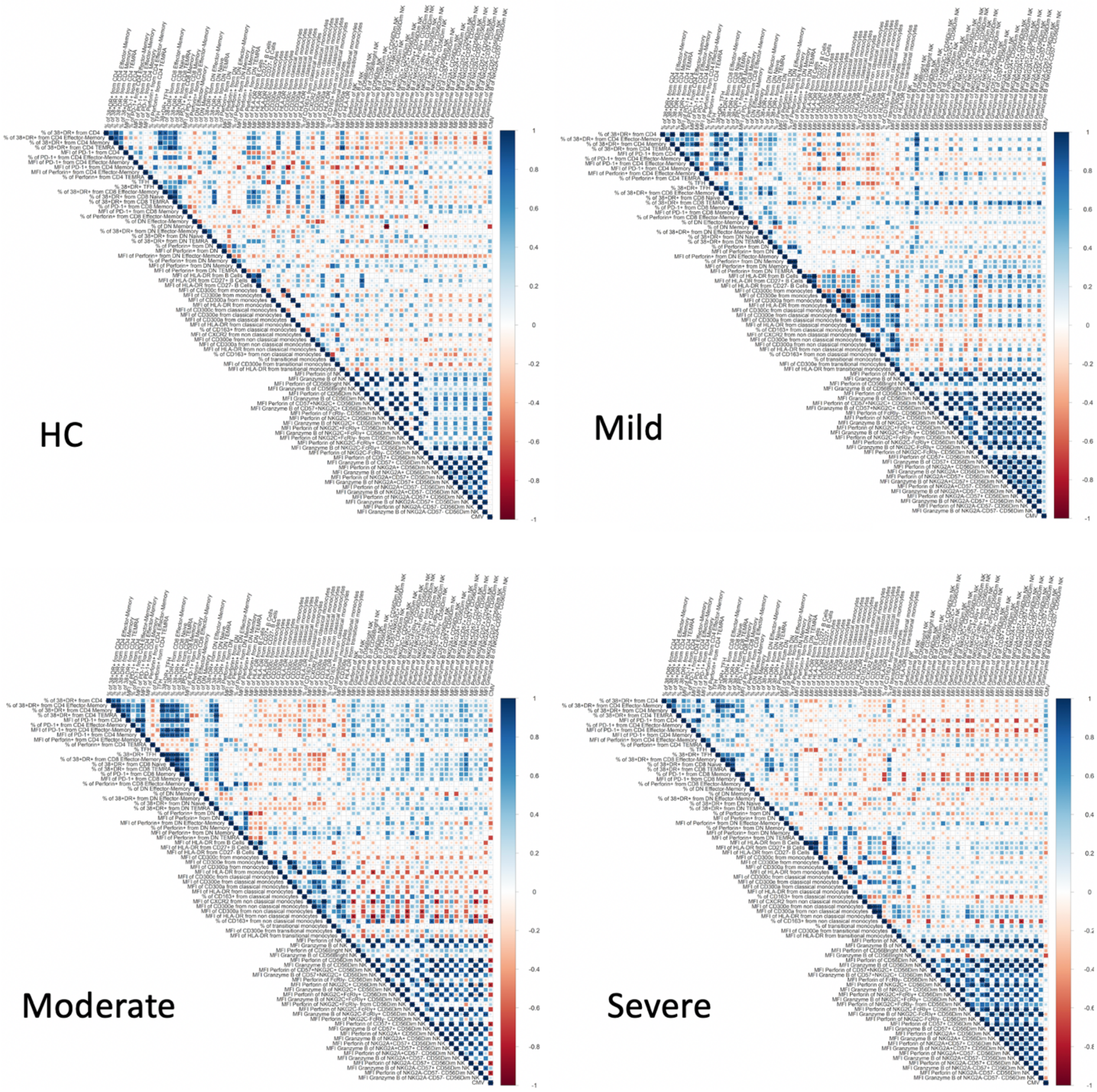

**Fig. S12. Correlation between significant flow cytometry variables.** Correlograms showing Spearman correlation of the significant flow cytometry variables for healthy controls (HC) and COVID-19 patients. *p <0.05, **p <0.001, and ***p <0.001.

**Table S1. Clinical characteristics of COVID-19 patients.**

| | | Mild | Moderate | Severe |
|---|---|---|---|---|
| **Group size** | n | 15 | 15 | 14 |
| **Risk Factors** | Age, median (range) | 62 (42-71) | 66 (40-83) | 68.5 (47-83) |
| | Sex, n (female/male) | 10/5 | 9/6 | 5/9 |
| | Smoking - Current, n (%) | 5 (33.33) | 1 (6.67) | 1 (7.14) |
| | - Prior, n (%) | 4 (26.67) | 3 (20) | 3 (21.43) |
| | - Non-smoker, n (%) | 4 (26.67) | 11 (73.33) | 5 (35.71) |
| | - Unknown, n (%) | 2 (13.33) | 0 | 5 (35.71) |
| **Comorbidities** | Hypertension, n (%) | 4 (26.67) | 2 (13.33) | 11 (78.57) |
| | Diabetes mellitus, n (%) | 3 (20) | 5 (33.33) | 3 (21.43) |
| | Coronary heart disease, n (%) | 1 (6.67) | 1 (6.67) | 0 |
| | Asthma, n (%) | 2 (13.33) | 3 (20) | 1 (7.14) |
| | Cancer, n (%) | 1 (6.67) | 1 (6.67) | 0 |
| | Others (COPD, obesity, autoinmune disease), n (%) | 4 (26.67) | 1 (6.67) | 2 (14.29) |
| | None, n (%) | 1 (6.67) | 2 (13.33) | 0 |
| **Symtomps on admission** | Fever, n (%) | 2 (13.33) | 11 (73.33) | 3 (21.43) |
| | Cough, n (%) | 7 (46.67) | 4 (26.67) | 10 (71.43) |
| | Dyspnea, n (%) | 10 (66.67) | 4 (26.67) | 7 (50) |
| | Body ache, n (%) | 4 (26.67) | 2 (13.33) | 2 (14.29) |
| | Days from symptom onset to study sampling, median (range) | 8 (0-35) | 3 (0-15) | 7 (0-21) |
| **Supportive oxygen therapy** | None, n (%) | 13 (86.67) | 7 (46.67) | 1 (7.14) |
| | <10 L/min, n (%) | 1 (6.67) | 4 (26.67) | 1 (7.14) |
| | 10-15 L/min, n (%) | 0 | 1 (6.67) | 1 (7.14) |
| | >15 L/min, n (%) | 0 | 2 (13.33) | 0 |
| | Ventilator, n (%) | 1 (6.67) | 1 (6.67) | 10 (71.43) |
| | ECMO, n (%) | 0 | 0 | 1 (7.14) |
| | Oxygen therapy, median in days (range) | 0 (0-28) | 1 (0-70) | 6 (0-16) |
| **Treatment prior to sampling, n (%)** | Antibiotics | 1 (6.67) | 1 (6.67) | 11 (78.57) |
| | Corticosteroids | 1 (6.67) | 0 | 2 (14.29) |
| | Antivirals | 1 (6.67) | 0 | 9 (64.29) |
| | Others | 0 | 2 (13.33) | 0 |
| **Clinical course** | Days on ICU, median (range) | 0 (0-10) | 0 (0-70) | 42 (0-79) |
| | Days intubated, median (range) | 0 (0-10) | 0 (0-53) | 21 (0-40) |
| | Days hospitalized, median (range) | 10 (0-31) | 11 (3-70) | 55 (13-115) |
| | Radiological bilateral infiltrations, n (%) | 4 (26.67) | 8 (53.33) | 12 (85.71) |
| | Embolism/thromosis | 0 | 0 | 4 (28.57) |
| **Outcome** | Discharged, n (%) | 15 (100) | 11 (73.33) | 12 (85.71) |
| | Death, n (%) | 0 | 4 (26.67) | 2 (14.29) |

**Table S2. Clinical laboratory results of COVID-19 patients.**

| | | Normal range | Mild | Moderate | Severe |
|---|---|---|---|---|---|
| | | | *Median (range)* | *Median (range)* | *Median (range)* |
| **At study sampling** | Leukocytes, cells*10^-3/µL | 4.5-11 | 8 (3.1-11.4) | 7.7 (4.6-17) | 11.35 (3.6-38.9) |
| | Neutrophils, cells*10^-3/µL | 2-5 | 4.1 (1.56-8.24) | 5.95 (3.39-16.03) | 9.84 (2.76-37.41) |
| | Monocytes, cells*10^-3/µL | 0-0.8 | 0.4 (0.15-1) | 0.41 (0.14-0.73) | 0.535 (0.30-1.06) |
| | Lymphocytes, cells*10^-3/µL | 1.3-2.9 | 1.77 (0.51-3.85) | 0.91 (0.26-2.08) | 0.93 (0.2-2.01) |
| | Neutrophils (%) | 43-65 | 60.50 (47.20-84.4) | 78 (59.8-94.2) | 84.15 (64.1-95.2) |
| | Monocytes (%) | 0-15 | 5.9 (1.7-9.2) | 5 (1.8-9) | 4.45 (1.9-9) |
| | Lymphocytes (%) | 20.5-45.5 | 30.3 (7.1-43.4) | 12 (2-30.8) | 8.5 (1.7-21.8) |
| | Platelets, cells*10^-3/µL | 135-450 | 250 (122-499) | 217 (111-301) | 303.5 (153-469) |
| | Hemoglobin, g/dL | 13-18 | 14 (12.4-16.9) | 13.3 (8.5-16.3) | 10.45 (7.6-14.8) |
| | C-reactive protein, mg/L | 0-11 | 1.94 (0.1-56.52) | 78.44 (1.16-225.48) | 47.31 (0.25-180) |
| | D-dimer, ng/mL | 0-500 | 450 (190-2790) | 650 (190-5320) | 2055 (760-7520) |
| | Ferritin, ng/mL | 15-300 | 189 (21-1958) | 261 (40-2922) | 840.5 (563-13010) |
| | Fibrinogen, mg/dL | 180-400 | 375 (274-750) | 750 (322-750) | 650.5 (139-750) |
| | Creatinine, mg/dL | 0.7-1.4 | 0.7 (0.36-0.97) | 0.85 (0.59-1.41) | 0.78 (0.32-2.75) |
| | IL-6, pg/L | 0-40 | 3 (3-8.3) | 37.1 (3-1139) | 12.6 (3-65.9) |
| **CMV Serology** | Positive, n (%) | | 11 (73.33) | 12 (80) | 10 (71.43) |
| | Negative, n (%) | | 3 (20) | 2 (13.33) | 3 (21.43) |
| **SARS-CoV-2 microbiological studies** | PCR +, n (%) | | 14 (93.33) | 15 (100) | 14 (100) |
| | PCR -, n (%) | | 1 (6.67) | 0 | 0 |
| | Antibodies - IgM, n (%) | | 2 (13.33) | 0 | 0 |
| | - IgG, n (%) | | 2 (13.33) | 3 (20) | 6 (42.86) |
| | - IgM, IgG, n (%) | | 3 (20) | 1 (6.67) | 6 (42.86) |
| | - Negative, n (%) | | 3 (20) | 8 (53.33) | 1 (7.14) |

**Table S3. Bivariate analysis of clinical laboratory and flow cytometry data.** Values are given as median and interquartile range, except gender and CMV status.

| | [ALL] | Mild | Moderate | Severe | N | p.value | p.adjus |
|---|---|---|---|---|---|---|---|
| **Gender: Male (%)** | 20 (45.5%) | 5 (33.3%) | 6 (40.0%) | 9 (64.3%) | 44 | 0.215 | 0.376 |
| **Gender: Female (%)** | 24 (54.5%) | 10 (66.7%) | 9 (60.0%) | 5 (35.7%) | | | |
| **Age** | 64.0 [55.0;70.2] | 62.0 [55.5;65.5] | 66.0 [53.5;73.0] | 68.5 [61.8;72.5] | 44 | 0.198 | 0.359 |
| **CMV: Negative, n (%)** | 8 (19.5%) | 3 (21.4%) | 2 (14.3%) | 3 (23.1%) | 41 | 0.892 | 0.929 |
| **CMV: Positive, n (%)** | 33 (80.5%) | 11 (78.6%) | 12 (85.7%) | 10 (76.9%) | | | |
| **Tryptase** | 4.10 [3.05;4.78] | 3.51 [2.96;4.61] | 4.56 [3.59;5.29] | 3.61 [2.52;4.57] | 42 | 0.193 | 0.353 |
| **IL-6** | 8.10 [3.00;42.1] | 3.00 [3.00;4.35] | 37.1 [13.1;136] | 12.6 [5.60;43.7] | 42 | <0.001 | 0.012 |
| **Monocyte count** | 0.42 [0.33;0.65] | 0.40 [0.23;0.62] | 0.41 [0.34;0.50] | 0.54 [0.44;0.84] | 44 | 0.059 | 0.193 |
| **% neutrophils** | 77.2 [64.0;85.7] | 60.5 [55.6;74.2] | 78.0 [71.8;88.8] | 84.2 [78.1;88.4] | 44 | <0.001 | 0.012 |
| **% lymphocytes** | 14.3 [7.33;24.4] | 30.3 [18.5;33.6] | 12.0 [5.45;20.1] | 8.50 [6.15;14.4] | 44 | <0.001 | 0.01 |
| **% monocytes** | 5.10 [4.07;6.73] | 5.90 [4.55;6.75] | 5.00 [3.60;6.10] | 4.45 [3.17;7.13] | 44 | 0.503 | 0.644 |
| **CRP** | 25.6 [2.34;77.9] | 1.94 [0.69;17.4] | 78.4 [20.9;157] | 47.3 [23.2;76.7] | 44 | <0.001 | 0.012 |
| **Leucocyte count** | 8.80 [6.42;11.5] | 8.00 [5.55;9.60] | 7.70 [6.20;11.2] | 11.4 [8.82;15.0] | 44 | 0.036 | 0.174 |
| **Neutrophil count** | 6.40 [4.01;9.76] | 4.19 [3.10;7.08] | 5.95 [4.34;10.4] | 9.84 [6.66;12.7] | 44 | 0.013 | 0.106 |
| **D-dimer** | 810 [412;2458] | 450 [240;660] | 650 [385;2715] | 2055 [1335;3995] | 44 | <0.001 | 0.012 |
| **Ferritin** | 507 [164;836] | 189 [73.0;352] | 261 [160;666] | 840 [716;1142] | 44 | <0.001 | 0.01 |
| **Fibrinogen** | 558 [405;750] | 375 [345;485] | 750 [516;750] | 650 [529;750] | 44 | 0.002 | 0.023 |
| **Creatinin** | 0.76 [0.66;0.90] | 0.70 [0.58;0.78] | 0.85 [0.70;1.02] | 0.78 [0.57;0.94] | 44 | 0.091 | 0.257 |
| **Hemoglobin** | 12.9 [10.8;14.2] | 14.0 [13.1;15.0] | 13.3 [11.7;14.4] | 10.4 [9.53;11.6] | 44 | <0.001 | 0.009 |
| **Lymphocyte count** | 1.21 [0.68;1.62] | 1.77 [1.29;2.11] | 0.91 [0.62;1.28] | 0.93 [0.44;1.23] | 44 | 0.005 | 0.05 |
| **Platelet count** | 250 [192;303] | 250 [200;313] | 217 [154;249] | 304 [259;399] | 44 | 0.003 | 0.031 |
| **% of CD4** | 70.0 [56.5;75.2] | 71.2 [63.3;75.0] | 71.0 [55.7;73.7] | 66.2 [51.1;78.1] | 44 | 0.853 | 0.912 |
| **% of CD8** | 25.7 [21.5;36.0] | 26.1 [21.3;32.8] | 23.6 [22.0;35.3] | 27.9 [19.7;44.0] | 44 | 0.7 | 0.787 |
| **% of DN** | 2.76 [1.71;4.17] | 2.02 [1.77;2.39] | 3.70 [3.12;6.14] | 2.84 [1.10;4.30] | 44 | 0.052 | 0.185 |
| **% of DP** | 0.40 [0.23;0.64] | 0.53 [0.40;0.74] | 0.33 [0.22;0.72] | 0.26 [0.17;0.43] | 44 | 0.05 | 0.18 |
| **CD4/CD8 Ratio** | 2.69 [1.54;3.47] | 2.73 [1.88;3.54] | 3.03 [1.52;3.31] | 2.44 [1.17;4.03] | 44 | 0.745 | 0.819 |
| **% of CD4** | 7.79 | 8.62 | 7.84 | 5.82 | 44 | 0.605 | 0.729 |

| | | | | | | | |
|---|---|---|---|---|---|---|---|
| **Effector-Memory** | [3.79;11.7] | [4.41;9.79] | [4.64;15.2] | [2.75;11.7] | | | |
| **% of CD4 Memory** | 48.0 [39.3;56.4] | 53.1 [48.3;60.2] | 45.9 [39.1;52.9] | 42.7 [36.3;50.6] | 44 | 0.108 | 0.286 |
| **% of CD4 Naive** | 36.9 [31.2;49.1] | 33.9 [31.8;44.7] | 37.2 [31.6;47.8] | 45.2 [32.5;54.9] | 44 | 0.418 | 0.56 |
| **% of CD4 TEMRA** | 0.39 [0.12;2.80] | 0.36 [0.09;2.55] | 0.43 [0.21;2.23] | 0.60 [0.09;3.96] | 44 | 0.746 | 0.819 |
| **% of 38+DR+ from CD4** | 2.20 [1.26;4.22] | 1.87 [0.94;2.52] | 1.97 [1.23;3.74] | 3.67 [1.49;6.59] | 44 | 0.151 | 0.326 |
| **% of 38+DR+ from CD4 Effector-Memory** | 5.78 [2.74;8.86] | 3.11 [1.31;7.32] | 4.47 [2.54;7.82] | 9.72 [6.39;17.2] | 44 | 0.023 | 0.139 |
| **% of 38+DR+ from CD4 Memory** | 3.13 [2.05;6.02] | 2.55 [1.36;3.93] | 3.19 [2.08;4.52] | 4.86 [2.52;13.0] | 44 | 0.137 | 0.318 |
| **% of 38+DR+ from CD4 Naive** | 0.15 [0.10;0.26] | 0.11 [0.09;0.22] | 0.15 [0.11;0.32] | 0.16 [0.12;0.27] | 44 | 0.607 | 0.729 |
| **% of 38+DR+ from CD4 TEMRA** | 4.09 [0.74;10.6] | 4.23 [1.09;7.73] | 1.81 [0.71;4.90] | 10.6 [0.20;15.8] | 44 | 0.263 | 0.401 |
| **% of PD-1+ from CD4** | 10.7 [7.55;15.1] | 7.88 [6.16;10.3] | 10.9 [8.67;13.8] | 14.7 [9.62;19.5] | 44 | 0.045 | 0.174 |
| **MFI of PD-1+ from CD4** | 1961 [1806;2101] | 1902 [1738;2012] | 1935 [1832;2058] | 2111 [1982;2494] | 44 | 0.022 | 0.139 |
| **% of PD-1+ from CD4 Effector-Memory** | 23.1 [17.2;40.4] | 15.8 [11.6;21.0] | 23.9 [19.4;36.8] | 37.1 [24.5;59.1] | 44 | 0.002 | 0.023 |
| **MFI of PD-1+ from CD4 Effector-Memory** | 1858 [1673;2210] | 1689 [1578;1872] | 1774 [1668;1970] | 2244 [1940;2590] | 44 | <0.001 | 0.013 |
| **% of PD-1+ from CD4 Memory** | 15.6 [12.0;22.7] | 12.4 [9.15;16.4] | 16.5 [13.5;23.6] | 20.6 [12.3;28.2] | 44 | 0.046 | 0.174 |
| **MFI of PD-1+ from CD4 Memory** | 2064 [1938;2324] | 1939 [1858;2108] | 2083 [1963;2156] | 2302 [2048;2568] | 44 | 0.021 | 0.139 |
| **% of PD-1+ from CD4 TEMRA** | 11.8 [5.30;22.2] | 11.3 [4.70;16.0] | 7.27 [5.12;23.4] | 16.5 [8.16;24.6] | 44 | 0.151 | 0.326 |
| **MFI of PD-1+ from CD4 TEMRA** | 1564 [1413;1822] | 1535 [1389;1980] | 1464 [1407;1564] | 1676 [1572;2309] | 41 | 0.146 | 0.326 |
| **% of Perforin+ from CD4** | 1280 [1132;1378] | 1325 [1262;1460] | 1213 [1099;1325] | 1249 [1140;1311] | 44 | 0.138 | 0.318 |
| **MFI of Perforin+ from CD4** | 1.79 [1.03;4.94] | 1.98 [1.08;5.40] | 1.81 [1.04;3.68] | 1.65 [0.90;5.80] | 44 | 0.844 | 0.907 |
| **% of Perforin+ from CD4 Effector-Memory** | 8.46 [2.13;15.8] | 9.77 [2.56;13.8] | 4.88 [2.10;13.9] | 7.32 [2.25;25.5] | 44 | 0.868 | 0.918 |
| **MFI of Perforin+ from CD4 Effector-Memory** | 1117 [1014;1275] | 1117 [1021;1285] | 1082 [956;1225] | 1130 [1028;1305] | 43 | 0.658 | 0.757 |
| **% of Perforin+ from CD4 Memory** | 0.82 [0.58;1.31] | 0.90 [0.68;1.29] | 0.68 [0.46;1.00] | 1.14 [0.52;1.70] | 44 | 0.294 | 0.429 |
| **MFI of Perforin+ from CD4 Memory** | 1170 [986;1304] | 1161 [970;1314] | 1168 [1005;1315] | 1203 [1006;1281] | 44 | 0.946 | 0.965 |
| **% of Perforin+ from CD4 TEMRA** | 37.8 [17.6;61.1] | 50.0 [19.6;64.7] | 28.1 [16.2;48.8] | 41.9 [19.0;57.7] | 44 | 0.668 | 0.758 |
| **MFI of Perforin+ from CD4 TEMRA** | 1791 [1288;4108] | 2087 [1537;2958] | 4024 [1218;5280] | 1410 [1292;3441] | 43 | 0.398 | 0.537 |

| | | | | | | | |
|---|---|---|---|---|---|---|---|
| **% TFH** | 6.86 [5.03;9.32] | 5.95 [4.76;8.10] | 7.73 [5.41;10.2] | 6.70 [4.95;9.44] | 44 | 0.524 | 0.652 |
| **% of CD8 Effector-Memory** | 11.8 [7.30;18.1] | 9.72 [7.87;16.5] | 9.73 [6.66;12.8] | 16.9 [12.0;33.4] | 44 | 0.053 | 0.185 |
| **% of CD8 Memory** | 32.3 [24.0;40.9] | 36.8 [25.8;44.9] | 31.9 [24.1;41.2] | 30.2 [20.0;37.4] | 44 | 0.421 | 0.56 |
| **% of CD8 Naive** | 15.8 [6.48;29.2] | 23.6 [10.2;31.6] | 15.2 [9.69;26.3] | 13.6 [4.42;22.2] | 44 | 0.217 | 0.376 |
| **% of CD8 TEMRA** | 15.5 [8.63;35.8] | 10.9 [6.53;23.0] | 18.8 [10.1;36.2] | 18.8 [13.8;38.1] | 44 | 0.38 | 0.515 |
| **% of 38+DR+ from CD8** | 9.87 [3.98;21.1] | 9.55 [2.22;13.2] | 9.09 [4.12;11.4] | 21.8 [10.1;52.6] | 44 | 0.034 | 0.174 |
| **% of 38+DR+ from CD8 Effector-Memory** | 13.1 [5.60;21.4] | 12.6 [3.39;18.8] | 10.5 [6.80;12.4] | 25.0 [18.0;71.5] | 44 | 0.014 | 0.109 |
| **% of 38+DR+ from CD8 Memory** | 14.6 [6.74;28.6] | 9.22 [3.14;24.4] | 10.6 [7.26;17.2] | 32.0 [17.0;64.0] | 44 | 0.018 | 0.127 |
| **% of 38+DR+ from CD8 Naive** | 0.87 [0.46;2.37] | 0.55 [0.30;1.11] | 0.64 [0.42;3.18] | 2.00 [0.87;5.17] | 44 | 0.037 | 0.174 |
| **% of 38+DR+ from CD8 TEMRA** | 7.77 [3.23;13.9] | 6.22 [2.74;10.0] | 8.13 [4.14;13.9] | 12.7 [3.48;37.1] | 44 | 0.215 | 0.376 |
| **% of PD-1+ from CD8** | 21.1 [14.8;33.4] | 16.7 [10.4;26.6] | 24.1 [18.9;36.6] | 22.4 [15.7;33.8] | 44 | 0.102 | 0.276 |
| **MFI of PD-1+ from CD8** | 1774 [1648;2022] | 1670 [1618;1826] | 1758 [1691;1987] | 1938 [1787;2154] | 44 | 0.043 | 0.174 |
| **% of PD-1+ from CD8 Effector-Memory** | 20.0 [10.9;35.9] | 12.2 [8.68;24.9] | 20.6 [15.7;42.0] | 21.2 [13.0;40.1] | 44 | 0.166 | 0.331 |
| **MFI of PD-1+ from CD8 Effector-Memory** | 1744 [1590;1920] | 1615 [1522;1760] | 1715 [1628;1982] | 1846 [1748;2117] | 44 | 0.039 | 0.174 |
| **% of PD-1+ from CD8 Memory** | 32.2 [22.7;43.4] | 22.8 [18.7;32.6] | 32.6 [29.4;41.4] | 41.2 [28.7;52.0] | 44 | 0.049 | 0.178 |
| **MFI of PD-1+ from CD8 Memory** | 2042 [1808;2233] | 1902 [1758;2026] | 2056 [1935;2314] | 2168 [1953;2286] | 44 | 0.037 | 0.174 |
| **% of PD-1+ from CD8 TEMRA** | 17.1 [9.72;30.2] | 10.0 [7.44;23.4] | 22.9 [17.4;32.3] | 15.9 [10.6;25.0] | 44 | 0.116 | 0.288 |
| **MFI of PD-1+ from CD8 TEMRA** | 1624 [1513;1730] | 1618 [1478;1648] | 1618 [1538;1709] | 1706 [1577;1744] | 44 | 0.222 | 0.379 |
| **% of Perforin+ from CD8** | 25.8 [16.3;46.7] | 24.4 [15.1;39.8] | 23.9 [12.8;36.4] | 42.9 [25.2;57.6] | 44 | 0.055 | 0.187 |
| **MFI of Perforin+ from CD8** | 1262 [1085;1477] | 1206 [1151;1366] | 1230 [979;1512] | 1328 [1106;1562] | 44 | 0.717 | 0.797 |
| **% of Perforin+ from CD8 Effector-Memory** | 46.9 [29.6;64.4] | 46.5 [30.6;64.8] | 43.0 [14.9;56.6] | 61.9 [39.4;68.8] | 44 | 0.116 | 0.288 |
| **MFI of Perforin+ from CD8 Effector-Memory** | 1266 [1060;1574] | 1220 [1151;1680] | 1244 [924;1506] | 1328 [1051;1560] | 44 | 0.606 | 0.729 |
| **% of Perforin+ from CD8 Memory** | 19.0 [11.3;27.9] | 18.7 [8.56;29.4] | 12.6 [10.7;20.5] | 25.6 [19.0;47.0] | 44 | 0.034 | 0.174 |
| **MFI of Perforin+ from CD8 Memory** | 984 [918;1111] | 976 [881;1050] | 1012 [944;1109] | 978 [932;1215] | 43 | 0.634 | 0.749 |
| **% of Perforin+ from CD8 TEMRA** | 63.2 [52.5;77.0] | 58.7 [53.1;76.2] | 69.7 [37.5;78.7] | 62.9 [60.7;72.9] | 44 | 0.881 | 0.927 |

| | | | | | | | |
|---|---|---|---|---|---|---|---|
| **MFI of Perforin+ from CD8 TEMRA** | 1470 [1273;1957] | 1639 [1382;2090] | 1415 [1150;2088] | 1512 [1234;1677] | 44 | 0.522 | 0.652 |
| **% of DN Effector-Memory** | 24.0 [18.6;33.8] | 21.5 [18.6;29.0] | 22.2 [17.4;34.5] | 29.0 [25.1;35.2] | 44 | 0.326 | 0.461 |
| **% of DN Memory** | 33.0 [20.3;46.2] | 40.4 [31.8;48.6] | 32.0 [18.4;40.3] | 20.0 [15.4;36.2] | 44 | 0.043 | 0.174 |
| **% of DN Naive** | 12.9 [5.79;20.6] | 12.0 [7.38;21.1] | 9.19 [4.26;16.6] | 15.0 [4.14;25.2] | 44 | 0.479 | 0.622 |
| **% of DN TEMRA** | 10.9 [4.74;19.2] | 8.16 [4.42;12.1] | 16.0 [9.30;19.2] | 9.38 [4.66;34.2] | 44 | 0.294 | 0.429 |
| **% of 38+DR+ from DN** | 6.88 [3.94;16.1] | 5.19 [3.27;9.99] | 7.37 [3.07;12.9] | 11.4 [6.82;23.0] | 44 | 0.064 | 0.205 |
| **% of 38+DR+ from DN Effector-Memory** | 3.18 [1.34;7.27] | 3.23 [1.04;8.92] | 3.13 [0.98;7.72] | 3.48 [1.48;5.94] | 44 | 0.978 | 0.983 |
| **% of 38+DR+ from DN Memory** | 6.52 [3.76;11.6] | 3.57 [2.32;8.25] | 5.20 [3.84;8.30] | 14.7 [5.91;39.4] | 44 | 0.027 | 0.154 |
| **% of 38+DR+ from DN Naive** | 18.7 [8.21;33.3] | 12.9 [7.16;18.7] | 22.4 [7.40;37.1] | 22.9 [15.6;42.7] | 44 | 0.059 | 0.193 |
| **% of 38+DR+ from DN TEMRA** | 8.27 [3.26;20.4] | 6.12 [2.50;13.0] | 6.38 [3.83;31.6] | 14.3 [5.96;21.3] | 44 | 0.258 | 0.4 |
| **% of Perforin+ from DN** | 51.0 [35.5;60.8] | 38.8 [31.0;57.5] | 51.4 [41.8;61.0] | 54.1 [48.3;63.9] | 44 | 0.111 | 0.287 |
| **MFI of Perforin+ from DN** | 2056 [1344;2957] | 1689 [1260;2297] | 2025 [1306;2804] | 2256 [1772;3553] | 44 | 0.188 | 0.35 |
| **% of Perforin+ from DN Effector-Memory** | 79.3 [62.4;87.8] | 66.7 [60.7;81.6] | 80.4 [61.5;88.2] | 83.4 [69.5;93.8] | 44 | 0.27 | 0.409 |
| **MFI of Perforin+ from DN Effector-Memory** | 4020 [2150;14491] | 2539 [2072;7508] | 3975 [2144;13535] | 9597 [4109;17924] | 44 | 0.285 | 0.425 |
| **% of Perforin+ from DN Memory** | 37.8 [28.8;48.8] | 32.3 [22.4;43.5] | 37.6 [30.8;44.8] | 44.5 [35.6;51.7] | 44 | 0.315 | 0.45 |
| **MFI of Perforin+ from DN Memory** | 1130 [1028;1240] | 1057 [988;1148] | 1143 [1046;1228] | 1226 [1077;1322] | 43 | 0.074 | 0.226 |
| **% of Perforin+ from DN TEMRA** | 78.3 [64.5;92.1] | 77.4 [68.8;87.8] | 84.4 [51.2;92.8] | 82.7 [71.7;93.7] | 44 | 0.863 | 0.918 |
| **MFI of Perforin+ from DN TEMRA** | 2588 [1882;3459] | 2676 [2272;3757] | 2702 [1273;3716] | 2522 [1882;2730] | 42 | 0.686 | 0.775 |
| **% of DP Effector-Memory** | 11.6 [6.36;18.9] | 11.4 [5.81;14.9] | 10.7 [8.44;18.9] | 13.4 [5.18;24.2] | 44 | 0.715 | 0.797 |
| **% of DP Memory** | 56.4 [42.7;60.9] | 58.6 [33.8;59.8] | 57.6 [45.8;61.3] | 55.3 [44.3;58.1] | 44 | 0.921 | 0.949 |
| **% of DP Naive** | 19.8 [11.2;28.6] | 19.0 [11.2;28.0] | 20.5 [17.2;28.6] | 19.6 [10.7;29.6] | 44 | 0.825 | 0.897 |
| **% of DP TEMRA** | 4.02 [1.27;16.4] | 4.47 [2.37;14.2] | 3.85 [2.12;18.4] | 5.18 [0.86;15.9] | 44 | 0.834 | 0.901 |
| **% of 38+DR+ from DP** | 5.52 [1.47;10.7] | 4.32 [1.32;12.9] | 6.06 [3.76;8.12] | 6.22 [1.23;10.2] | 44 | 0.987 | 0.987 |
| **% of 38+DR+ from DP Effector-Memory** | 1.53 [0.00;9.30] | 4.17 [1.69;16.0] | 0.00 [0.00;2.50] | 0.00 [0.00;10.5] | 44 | 0.023 | 0.139 |
| **% of 38+DR+ from DP Memory** | 4.93 [1.01;12.6] | 4.31 [1.21;16.6] | 5.56 [1.48;9.82] | 5.18 [0.74;12.0] | 44 | 0.889 | 0.929 |
| **% of 38+DR+ from DP Naive** | 0.00 [0.00;3.82] | 1.72 [0.00;13.7] | 0.75 [0.00;3.20] | 0.00 [0.00;0.00] | 44 | 0.161 | 0.329 |
| **% of 38+DR+** | 0.00 | 6.90 | 0.00 | 0.00 | 44 | 0.478 | 0.622 |

| | | | | | | | |
|---|---|---|---|---|---|---|---|
| **from DP TEMRA** | [0.00;12.8] | [0.00;11.9] | [0.00;12.9] | [0.00;15.0] | | | |
| **% of Perforin+ from DP** | 9.16 [3.63;29.2] | 19.6 [5.68;35.3] | 4.94 [0.65;13.3] | 10.9 [5.83;25.4] | 44 | 0.169 | 0.332 |
| **MFI of Perforin+ from DP** | 1335 [1116;1686] | 1360 [1088;1443] | 1304 [1178;2317] | 1340 [1095;1877] | 40 | 0.62 | 0.737 |
| **% of Perforin+ from DP Effector-Memory** | 8.39 [0.00;21.9] | 12.8 [3.69;23.1] | 0.00 [0.00;9.54] | 17.4 [0.00;24.7] | 44 | 0.045 | 0.174 |
| **MFI of Perforin+ from DP Effector-Memory** | 1512 [1080;2164] | 1562 [1358;1718] | 1161 [1046;1700] | 1340 [1065;2500] | 29 | 0.496 | 0.639 |
| **% of Perforin+ from DP Memory** | 5.07 [0.32;13.8] | 7.32 [4.78;14.8] | 3.65 [0.00;5.99] | 5.32 [0.00;17.7] | 44 | 0.06 | 0.195 |
| **MFI of Perforin+ from DP Memory** | 1188 [1030;1371] | 1136 [980;1366] | 1141 [1046;1213] | 1275 [1239;2033] | 33 | 0.07 | 0.217 |
| **% of Perforin+ from DP TEMRA** | 31.2 [0.00;69.2] | 53.4 [0.00;76.0] | 0.00 [0.00;74.4] | 26.8 [0.00;55.3] | 44 | 0.606 | 0.729 |
| **MFI of Perforin+ from DP TEMRA** | 1426 [1261;1709] | 1426 [1392;1484] | 1751 [1192;3498] | 1320 [1165;1518] | 26 | 0.663 | 0.757 |
| **MFI of HLA-DR from B Cells** | 11136 [7798;12810] | 11963 [8808;13441] | 11507 [10074;12906] | 9186 [6528;12524] | 44 | 0.54 | 0.666 |
| **MFI of CXCR5 from B Cells** | 3296 [2850;3841] | 3746 [2955;3866] | 3089 [2860;3635] | 3268 [2731;3692] | 44 | 0.448 | 0.592 |
| **% of CD27+ from B cells** | 14.4 [9.27;31.2] | 20.3 [8.83;40.8] | 13.9 [10.8;26.4] | 10.1 [8.03;19.3] | 44 | 0.257 | 0.4 |
| **MFI of HLA-DR from CD27+ B Cells** | 8383 [6943;9952] | 8524 [6952;10042] | 9406 [8469;10935] | 7488 [5259;8314] | 44 | 0.028 | 0.157 |
| **MFI of CXCR5 from CD27+ B Cells** | 2914 [2507;3154] | 3095 [2806;3513] | 2843 [2486;3117] | 2707 [2150;3036] | 44 | 0.163 | 0.329 |
| **% of Plasmablast from CD27+ B cells** | 0.68 [0.27;1.19] | 0.52 [0.29;1.24] | 1.07 [0.78;1.64] | 0.41 [0.02;0.69] | 44 | 0.016 | 0.121 |
| **% of CD27- from B cells** | 85.7 [68.8;90.7] | 79.7 [59.2;91.2] | 86.1 [73.6;89.2] | 89.8 [80.7;92.0] | 44 | 0.263 | 0.401 |
| **MFI of HLA-DR from CD27- B Cells** | 12256 [7934;14943] | 12101 [9524;15483] | 12698 [10192;15646] | 10020 [5426;13688] | 44 | 0.507 | 0.646 |
| **MFI of CXCR5 from CD27- B Cells** | 3406 [2950;4003] | 3878 [3172;4028] | 3234 [2937;3800] | 3304 [2775;3833] | 44 | 0.375 | 0.512 |
| **MFI of CD300c from monocytes** | 996 [587;1606] | 1429 [806;2077] | 626 [408;1350] | 1069 [606;1529] | 41 | 0.201 | 0.36 |
| **MFI of CXCR2 from monocytes** | 9115 [8045;10699] | 9282 [8045;10490] | 8768 [7721;11193] | 9343 [8355;10133] | 41 | 0.954 | 0.968 |
| **MFI of CD300e from monocytes** | 1686 [1181;2193] | 1896 [1394;2190] | 2225 [1764;2707] | 945 [594;1230] | 41 | <0.001 | 0.008 |
| **MFI of CD300a from monocytes** | 2877 [2485;4831] | 2802 [2605;3332] | 5084 [3667;6081] | 2132 [1877;2837] | 41 | 0.004 | 0.039 |
| **MFI of HLA-DR from monocytes** | 2529 [1815;4091] | 3946 [2140;5410] | 2748 [1962;3634] | 2355 [1283;2639] | 41 | 0.153 | 0.326 |
| **% of CD163+ from monocytes** | 10.7 [2.60;22.3] | 2.91 [1.49;9.69] | 20.2 [5.06;25.7] | 11.9 [7.45;17.2] | 41 | 0.032 | 0.171 |
| **% of classical monocytes** | 91.3 [87.1;92.4] | 91.3 [88.6;94.3] | 88.9 [85.6;91.5] | 92.1 [87.1;92.6] | 41 | 0.203 | 0.361 |
| **MFI of CD300c from classical** | 1004 [594;1674] | 1427 [820;2097] | 638 [408;1342] | 1072 [625;1615] | 41 | 0.171 | 0.334 |

| | | | | | | | |
|---|---|---|---|---|---|---|---|
| **monocytes** | | | | | | | |
| **MFI of CXCR2 from classical monocytes** | 9491 [8448;11377] | 9545 [8337;10828] | 9262 [8001;11586] | 9929 [8620;10846] | 41 | 0.935 | 0.959 |
| **MFI of CD300e from classical monocytes** | 1509 [1119;2103] | 1852 [1338;2101] | 2014 [1594;2409] | 840 [571;1121] | 41 | <0.001 | 0.008 |
| **MFI of CD300a from classical monocytes** | 2771 [2419;4701] | 2695 [2482;3253] | 4886 [3563;5631] | 2083 [1846;2681] | 41 | 0.003 | 0.031 |
| **MFI of HLA-DR from classical monocytes** | 2301 [1709;3847] | 3634 [1956;4918] | 2096 [1738;3290] | 1972 [1249;2471] | 41 | 0.132 | 0.311 |
| **% of CD163+ from classical monocytes** | 8.31 [1.96;16.3] | 2.46 [1.07;9.58] | 14.3 [4.23;23.1] | 10.2 [5.30;13.8] | 41 | 0.046 | 0.174 |
| **% of non classical monocytes** | 1.75 [1.01;2.81] | 2.04 [1.17;3.66] | 1.56 [1.04;2.13] | 1.60 [0.81;2.81] | 41 | 0.643 | 0.755 |
| **MFI of CD300c from non classical monocytes** | 777 [518;1172] | 993 [560;1310] | 579 [339;716] | 860 [663;1541] | 41 | 0.127 | 0.307 |
| **MFI of CXCR2 from non classical monocytes** | 419 [310;679] | 477 [409;711] | 425 [336;663] | 367 [207;407] | 41 | 0.101 | 0.276 |
| **MFI of CD300e from non classical monocytes** | 2514 [1014;3046] | 2568 [1858;2816] | 3590 [2564;4386] | 521 [263;1987] | 41 | 0.001 | 0.017 |
| **MFI of CD300a from non classical monocytes** | 8135 [1822;11095] | 8996 [7710;10912] | 11087 [8501;14143] | 1702 [1089;5302] | 41 | <0.001 | 0.01 |
| **MFI of HLA-DR from non classical monocytes** | 9996 [2144;14822] | 13490 [7425;18297] | 11561 [9665;18254] | 1881 [1008;4761] | 41 | 0.002 | 0.025 |
| **% of CD163+ from non classical monocytes** | 9.79 [4.37;21.9] | 4.03 [2.20;14.8] | 10.8 [6.54;21.3] | 16.4 [6.58;31.2] | 41 | 0.012 | 0.103 |
| **% of transitional monocytes** | 6.02 [4.19;9.80] | 4.42 [2.99;8.04] | 7.58 [6.19;11.5] | 5.90 [4.32;9.26] | 41 | 0.044 | 0.174 |
| **MFI of CD300c from trasitional monocytes** | 747 [447;1325] | 680 [527;1307] | 624 [386;1144] | 831 [483;1340] | 41 | 0.526 | 0.652 |
| **MFI of CXCR2 from trasitional monocytes** | 3991 [2461;6054] | 3286 [2212;4672] | 4016 [2588;5213] | 4711 [3671;7596] | 41 | 0.254 | 0.4 |
| **MFI of CD300e from trasitional monocytes** | 3590 [2782;4913] | 3458 [2822;4012] | 5600 [4024;5989] | 2670 [1964;3279] | 41 | 0.002 | 0.023 |
| **MFI of CD300a from trasitional monocytes** | 9280 [5592;11666] | 9450 [7052;11267] | 11774 [9475;13004] | 5701 [4416;7221] | 41 | 0.023 | 0.139 |
| **MFI of HLA-DR from trasitional monocytes** | 17408 [9751;28410] | 26764 [15962;29343] | 17820 [13536;27348] | 9751 [6120;17408] | 41 | 0.075 | 0.226 |
| **% of CD163+ from trasitional monocytes** | 20.9 [8.40;55.3] | 10.4 [5.71;19.5] | 42.3 [12.9;60.8] | 42.3 [19.6;57.7] | 41 | 0.047 | 0.174 |
| **MFI of CD300a from granulocytes** | 1174 [762;1484] | 1190 [864;1350] | 1483 [1179;1548] | 755 [658;1074] | 41 | 0.008 | 0.074 |
| **% of CD56Bright** | 3.82 | 4.91 | 2.60 | 2.46 | 43 | 0.232 | 0.386 |

| | | | | | | | |
|---|---|---|---|---|---|---|---|
| **NK** | [1.14;5.43] | [2.67;5.63] | [1.38;5.59] | [0.59;4.27] | | | |
| **% of CD56Dim NK** | 91.4 [88.1;94.2] | 91.4 [88.5;93.0] | 91.7 [90.3;95.4] | 90.1 [86.1;93.8] | 43 | 0.609 | 0.729 |
| **% of CD57+NKG2C+ from CD56Dim NK** | 6.79 [2.56;14.1] | 4.38 [1.54;10.9] | 12.4 [6.20;19.0] | 5.36 [3.81;8.58] | 43 | 0.077 | 0.226 |
| **% of FcRIγ- from CD56Dim NK** | 3.53 [1.23;15.2] | 2.07 [0.98;7.96] | 7.73 [2.42;27.4] | 3.37 [2.53;15.8] | 43 | 0.181 | 0.346 |
| **% of NKG2C+ from CD56Dim NK** | 8.64 [4.12;18.0] | 6.15 [2.84;13.4] | 17.8 [12.9;25.4] | 7.04 [4.62;11.2] | 43 | 0.022 | 0.139 |
| **% of NKG2C+FcRIγ+ from CD56Dim NK** | 4.25 [2.25;10.1] | 3.00 [1.77;5.04] | 9.61 [2.65;13.1] | 3.95 [3.03;9.57] | 43 | 0.153 | 0.326 |
| **% of NKG2C+FcRIγ- from CD56Dim NK** | 0.54 [0.04;3.80] | 0.10 [0.02;1.67] | 2.68 [0.61;4.34] | 0.17 [0.02;2.11] | 43 | 0.103 | 0.276 |
| **% of NKG2C-FcRIγ+ from CD56Dim NK** | 84.5 [69.1;93.1] | 91.6 [82.2;95.1] | 71.0 [55.8;84.5] | 84.8 [73.8;91.6] | 43 | 0.011 | 0.093 |
| **% of NKG2C-FcRIγ- from CD56Dim NK** | 3.27 [0.84;9.14] | 1.60 [0.70;4.56] | 4.89 [1.20;12.1] | 3.76 [2.51;9.79] | 43 | 0.19 | 0.35 |
| **% of CD56- NK** | 2.69 [1.10;4.23] | 3.37 [1.23;4.18] | 1.29 [1.00;3.92] | 3.05 [1.20;6.04] | 43 | 0.312 | 0.45 |
| **MFI Perforin of NK** | 6972 [5246;10184] | 8775 [6428;10208] | 6475 [4607;8150] | 6324 [5246;10584] | 43 | 0.349 | 0.491 |
| **MFI Granzyme B of NK** | 7301 [5360;8974] | 5799 [4060;7999] | 7530 [6153;8682] | 7431 [6308;10889] | 43 | 0.154 | 0.326 |
| **MFI Perforin of CD56Bright NK** | 1777 [1268;2257] | 1784 [1370;2285] | 1564 [1250;1954] | 1997 [1726;2558] | 43 | 0.156 | 0.327 |
| **MFI Granzyme B of CD56Bright NK** | 1658 [935;3931] | 1246 [366;1832] | 1149 [785;1942] | 3863 [2778;6837] | 43 | <0.001 | 0.015 |
| **MFI Perforin of CD56Dim NK** | 7512 [5976;10870] | 9156 [7093;10742] | 6870 [4630;8554] | 7124 [5957;10976] | 43 | 0.258 | 0.4 |
| **MFI Granzyme B of CD56Dim NK** | 7496 [5472;9652] | 6160 [4040;8418] | 7936 [7146;8687] | 8642 [6272;11229] | 43 | 0.152 | 0.326 |
| **MFI Perforin of CD57+NKG2C+ CD56Dim NK** | 8600 [5205;12170] | 10318 [6958;14086] | 6334 [4926;9752] | 8278 [6696;11533] | 43 | 0.233 | 0.386 |
| **MFI NKG2C of CD57+NKG2C+ CD56Dim NK** | 1597 [1512;1844] | 1521 [1454;1772] | 1636 [1559;1903] | 1628 [1554;1718] | 43 | 0.247 | 0.397 |
| **MFI Granzyme B of CD57+NKG2C+ CD56Dim NK** | 11455 [7344;13055] | 9385 [6518;12034] | 11468 [8562;12460] | 11888 [9417;17276] | 43 | 0.126 | 0.307 |
| **MFI Perforin of FcRIγ- CD56Dim NK** | 3561 [1814;5782] | 3083 [1624;4788] | 4449 [3192;5924] | 3154 [2707;6456] | 43 | 0.662 | 0.757 |
| **MFI Granzyme B of FcRIγ- CD56Dim NK** | 6173 [2212;9014] | 2629 [514;5792] | 8574 [6443;10421] | 6788 [4054;9325] | 43 | 0.078 | 0.226 |
| **MFI Perforin of NKG2C+ CD56Dim NK** | 7957 [5100;10772] | 9906 [6198;13106] | 6297 [4509;9616] | 8184 [5819;10669] | 43 | 0.258 | 0.4 |
| **MFI NKG2C of** | 1529 | 1489 | 1592 | 1528 | 43 | 0.356 | 0.492 |

| | | | | | | | |
|---|---|---|---|---|---|---|---|
| **NKG2C+ CD56Dim NK** | [1466;1774] | [1413;1657] | [1489;1872] | [1490;1717] | | | |
| **MFI Granzyme B of NKG2C+ CD56Dim NK** | 10602 [7020;12504] | 7886 [5784;11148] | 11146 [8169;11814] | 11450 [9075;16672] | 43 | 0.045 | 0.174 |
| **MFI Perforin of NKG2C+FcRIγ+ CD56Dim NK** | 8736 [5654;11827] | 9974 [7049;13159] | 7390 [4445;11314] | 9008 [6566;11091] | 43 | 0.479 | 0.622 |
| **MFI NKG2C of NKG2C+FcRIγ+ CD56Dim NK** | 1670 [1612;1926] | 1633 [1536;1829] | 1826 [1651;1965] | 1661 [1616;1857] | 43 | 0.141 | 0.322 |
| **MFI Granzyme B of NKG2C+FcRIγ+ CD56Dim NK** | 10970 [7373;12563] | 8854 [6522;11380] | 10995 [7928;11456] | 12400 [9162;17091] | 43 | 0.039 | 0.174 |
| **MFI Perforin of NKG2C+FcRIγ- from CD56Dim NK** | 4159 [2754;6440] | 4430 [2308;7156] | 4159 [2558;6240] | 4185 [3028;6430] | 35 | 0.972 | 0.982 |
| **MFI NKG2C of NKG2C+FcRIγ- from CD56Dim NK** | 1702 [1592;2096] | 1720 [1602;2326] | 1794 [1621;1928] | 1621 [1467;1696] | 35 | 0.315 | 0.45 |
| **MFI Granzyme B of NKG2C+FcRIγ- from CD56Dim NK** | 9642 [6028;11983] | 7356 [4507;11702] | 10294 [8208;11613] | 10074 [7944;15098] | 35 | 0.357 | 0.492 |
| **MFI Perforin of NKG2C-FcRIγ+ CD56Dim NK** | 8355 [6755;11058] | 9301 [7544;12188] | 7622 [6910;9064] | 7654 [6294;11064] | 43 | 0.356 | 0.492 |
| **MFI Granzyme B of NKG2C-FcRIγ+ CD56Dim NK** | 6911 [4940;8534] | 6160 [4028;7585] | 6650 [5414;7430] | 7928 [6058;10824] | 43 | 0.111 | 0.287 |
| **MFI Perforin of NKG2C-FcRIγ- CD56Dim NK** | 3561 [2070;5462] | 2849 [1658;4832] | 4562 [3590;5492] | 3407 [2648;6386] | 43 | 0.662 | 0.757 |
| **MFI Granzyme B of NKG2C-FcRIγ- CD56Dim NK** | 5925 [2488;9958] | 3004 [512;5925] | 8411 [5216;10590] | 6440 [3564;9277] | 43 | 0.067 | 0.21 |
| **% of CD57+ from CD56Bright NK** | 3.33 [0.86;6.53] | 2.37 [1.24;3.79] | 2.45 [0.62;6.57] | 5.61 [1.22;13.8] | 43 | 0.236 | 0.388 |
| **% of NKG2A+ from CD56Bright NK** | 91.9 [86.3;95.3] | 91.1 [90.2;93.4] | 94.9 [91.7;96.6] | 87.8 [76.1;96.2] | 43 | 0.182 | 0.346 |
| **% of CD57+ from CD56Dim NK** | 46.6 [31.8;60.0] | 36.1 [30.3;54.0] | 56.0 [40.3;67.3] | 49.4 [31.6;59.6] | 43 | 0.113 | 0.288 |
| **MFI Perforin of CD57+ CD56Dim NK** | 8854 [6390;11653] | 10435 [7752;12518] | 7138 [6062;9685] | 7589 [5824;11601] | 43 | 0.226 | 0.382 |
| **MFI Granzyme B of CD57+ CD56Dim NK** | 8794 [6166;10821] | 7748 [4975;9673] | 9455 [7940;10596] | 9640 [7878;13500] | 43 | 0.279 | 0.418 |
| **% of NKG2A+ from CD56Dim NK** | 36.8 [27.4;44.1] | 35.8 [29.5;43.0] | 34.8 [21.0;44.6] | 39.9 [30.8;42.6] | 43 | 0.918 | 0.949 |
| **MFI Perforin of NKG2A+ CD56Dim NK** | 7221 [5626;9862] | 8117 [5970;10677] | 6667 [5629;8304] | 6676 [5428;10199] | 43 | 0.661 | 0.757 |
| **MFI Granzyme B of NKG2A+** | 7957 [5247;9556] | 5912 [3852;8936] | 7280 [6103;8869] | 9530 [6066;11673] | 43 | 0.089 | 0.256 |

| | | | | | | | |
|---|---|---|---|---|---|---|---|
| **CD56Dim NK** | | | | | | | |
| **% of NKG2A+CD57+ from CD56Dim NK** | 12.3 [7.18;17.4] | 10.1 [7.08;17.0] | 13.9 [5.92;21.6] | 12.2 [8.39;15.6] | 43 | 0.764 | 0.834 |
| **MFI Perforin of NKG2A+CD57+ CD56Dim NK** | 9449 [6806;11894] | 10747 [8819;12402] | 8177 [6118;10691] | 8136 [6706;12000] | 43 | 0.246 | 0.397 |
| **MFI Granzyme B of NKG2A+CD57+ CD56Dim NK** | 9034 [6638;11455] | 7992 [4997;10284] | 8883 [6816;11312] | 11377 [7852;12941] | 43 | 0.178 | 0.345 |
| **% of NKG2A+CD57- from CD56Dim NK** | 22.4 [16.6;27.6] | 22.8 [18.2;29.6] | 15.8 [12.2;25.7] | 24.2 [19.9;28.1] | 43 | 0.101 | 0.276 |
| **MFI Perforin of NKG2A+CD57- CD56Dim NK** | 6516 [4938;8299] | 6881 [4938;9178] | 5956 [5454;7006] | 6428 [4664;8602] | 43 | 0.513 | 0.649 |
| **MFI Granzyme B of NKG2A+CD57- CD56Dim NK** | 6602 [4554;8162] | 4944 [3400;7714] | 5843 [5213;7512] | 8620 [5672;11103] | 43 | 0.042 | 0.174 |
| **% of NKG2A-CD57+ from CD56Dim NK** | 28.8 [20.4;37.9] | 25.7 [19.7;30.9] | 33.0 [20.2;47.1] | 32.8 [21.4;39.5] | 43 | 0.295 | 0.429 |
| **MFI Perforin of NKG2A-CD57+ CD56Dim NK** | 8543 [6120;11693] | 10248 [7490;12634] | 7084 [5986;9762] | 7692 [5743;11707] | 43 | 0.219 | 0.376 |
| **MFI Granzyme B of NKG2A-CD57+ CD56Dim NK** | 8677 [6068;10945] | 7646 [5060;9454] | 9444 [8304;10852] | 9456 [7723;13782] | 43 | 0.189 | 0.35 |
| **% of NKG2A-CD57- from CD56Dim NK** | 26.4 [18.6;41.8] | 35.7 [24.9;46.5] | 22.6 [15.4;33.7] | 23.0 [18.7;35.8] | 43 | 0.164 | 0.329 |
| **MFI Perforin of NKG2A-CD57- CD56Dim NK** | 8024 (3210) | 9043 (3789) | 6679 (2990) | 8276 (2384) | 43 | 0.131 | 0.311 |
| **MFI Granzyme B of NKG2A-CD57- CD56Dim NK** | 6296 (2580) | 5655 (2973) | 6051 (1262) | 7226 (2988) | 43 | 0.243 | 0.396 |
| **% 38+DR+ TFH** | 7.13 [4.09;13.9] | 5.14 [3.55;14.4] | 6.12 [4.42;8.72] | 11.3 [6.76;16.7] | 44 | 0.159 | 0.329 |

### Table S4. Flow cytometry panels.

#### T and B LYMPHOCYTES

| Laser | Filter | Fluorochrome | Marker | Manufacturer | Clone |
|---|---|---|---|---|---|
| **405** | 450/50 | BV421 | Perforin | BD Biosciences | δg9 |
| | 525/50 | BV510 | CD3 | BD Biosciences | UCTH1 |
| | 610/20 | BV605 | HLA-DR | BD Biosciences | G46-6 |
| | 670/30 | BV650 | CD8 | BD Biosciences | RPA-T8 |
| | 710/50 | BV711 | PD-1 | BD Biosciences | EH12.1 |
| | 780/60 | BV786 | CD38 | BD Biosciences | HIT2 |
| **488** | 530/30 | FITC | CD4 | BioLegend | RPA-T4 |
| | 710/50 | PercP-Cy5.5 | CD45RA | eBioscience | HI100 |
| | 780/60 | PE-Cy7 | CXCR5 | eBioscience | MU5UBEE |
| **640** | 670/30 | APC | CD27 | BioLegend | O323 |
| | 730/45 | AF700 | CD19 | eBioscience | SJ25C1 |

#### NK CELLS

| Laser | Filter | Fluorochrome | Marker | Manufacturer | Clone |
|---|---|---|---|---|---|
| **405** | 450/50 | BV421 | Perforin | BD Biosciences | δg9 |
| | 525/50 | BV510 | CD3/CD14/CD19 | BD Biosciences | UCHT1/MφP9/SJ25C1 |
| | 610/20 | BV605 | CD16 | BD Biosciences | 3G8 |
| | 710/50 | BV711 | CD56 | BD Biosciences | NCAM16.2 |
| | 780/60 | BV786 | NKG2C | BD Biosciences | 134591 |
| **488** | 530/30 | FITC | FcεRIγ | Milli-Mark | Polyclonal |
| | 575/25 | PE | Granzyme B | BD Biosciences | GB11 |
| | 780/60 | PE-Cy7 | NKp80 | Miltenyi Biotec | 4A4.D10 |
| **640** | 670/30 | APC | NKG2A | Beckman Coulter | Z199 |
| | 780/60 | APC-Cy7 | CD57 | Miltenyi Biotec | REA769 |

#### MONOCYTES

| Laser | Filter | Fluorochrome | Marker | Manufacturer | Clone |
|---|---|---|---|---|---|
| **405** | 610/20 | BV605 | CD163 | BioLegend | GHI/61 |
| | 710/50 | BV711 | CCR2 | BioLegend | K036C2 |
| | 780/60 | BV786 | CD300e | BD Biosciences | UP-H1 |
| **488** | 530/30 | FITC | CD16 | BD Biosciences | 3G8 |
| | 575/25 | PE | CD300a | Beckman Coulter | E59.126 |
| | 710/50 | PercP-Cy5.5 | HLA-DR | BD Biosciences | G46-6 |
| | 780/60 | PE-Cy7 | CD14 | BD Biosciences | MφP9 |
| **640** | 670/30 | APC | CD300c | eBioscience | TX45 |
| | 780/60 | APC-Cy7 | CD66b | BioLegend | G10F5 |
